# Endoplasmic reticulum oxidoreductin (ERO) provides resilience against reductive stress and hypoxic conditions by mediating luminal redox dynamics

**DOI:** 10.1101/2021.12.13.472397

**Authors:** José Manuel Ugalde, Isabel Aller, Lika Kudrjasova, Romy R. Schmidt, Michelle Schlößer, Maria Homagk, Philippe Fuchs, Sophie Lichtenauer, Markus Schwarzländer, Stefanie J. Müller-Schüssele, Andreas J. Meyer

**Affiliations:** INRES-Chemical Signalling, University of Bonn, Friedrich-Ebert-Allee 144, D-53113 Bonn, Germany; Plant Biotechnology, Bielefeld University, D-33615 Bielefeld, Germany; Institute for Biology and Biotechnology of Plants, University of Münster, Schlossplatz 8, D- 48143 Münster, Germany; Molecular Botany, Department of Biology, TU Kaiserslautern, Erwin-Schrödinger-Straße 70, D-67663, Kaiserslautern, Germany

**Keywords:** Endoplasmic reticulum, oxidative protein folding, ER oxidoreductin, glutathione redox potential, redox-sensitive GFP, redox imaging, unfolded protein response, hypoxia

## Abstract

Oxidative protein folding in the endoplasmic reticulum (ER) depends on the coordinated action of protein disulfide isomerases and ER oxidoreductins (EROs). Strict dependence of ERO activity on molecular oxygen as the final electron acceptor implies that oxidative protein folding and other ER processes are severely compromised under hypoxia. While many key players involved in oxidative protein folding are known, our understanding of how redox homeostasis in the ER is maintained and how EROs, the Cys residues of nascent proteins, and the luminal glutathione redox buffer interact is limited. Here, we isolated viable *ero1 ero2* double mutants largely deficient in ERO activity, which rendered the mutants highly sensitive to reductive stress and hypoxia. To elucidate the specific redox dynamics in the ER lumen *in vivo*, we expressed the glutathione redox potential (*E*_GSH_) sensor Grx1-roGFP2iL-HDEL with a midpoint potential of -240 mV in the ER of Arabidopsis plants. We found *E*_GSH_ values of -241 mV in wild-type plants, which is less oxidizing than previously estimated. In the *ero1 ero2* mutants, luminal *E*_GSH_ was reduced further to -253 mV. Recovery to reductive ER stress, as induced by acute exposure to dithiothreitol, was delayed in *ero1 ero2* mutants. The characteristic signature of *E*_GSH_ dynamics in the ER lumen triggered by hypoxia was affected in the *ero1 ero2* mutant reflecting a disrupted balance of reductive and oxidizing inputs, including nascent polypeptides and glutathione entry. The ER redox dynamics can now be dissected *in vivo*, revealing a central role of EROs as major redox integrators to promote luminal redox homeostasis.

**One sentence summary:** Dynamic monitoring the ER luminal glutathione redox potential highlights the role of EROs in defining redox conditions and the interplay between different redox inputs during hypoxia and reductive stress.

## INTRODUCTION

Many proteins rely on the formation of disulfide bonds as crucial post-translational modifications for their structure and function. Those disulfides are generated by distinct oxidative protein folding mechanisms that operate in different subcellular locations (Meyer et al., 2019). The endoplasmic reticulum (ER) is the primary cellular site of oxidative protein folding, supplying the continuous flux of proteins that pass through the secretory pathway with disulfide bonds. Under standard conditions *de novo* formation of disulfide bonds in the ER lumen is achieved by transferring electrons from cysteines in nascent peptides to molecular oxygen as the final electron acceptor. The electron transfer is mediated by a disulfide relay system consisting of members of the protein disulfide isomerase (PDI) family and thiol oxidases named endoplasmic reticulum oxidoreductins (EROs) (Matsusaki et al., 2016; Fan et al., 2019). In the first step of this electron transfer cascade, oxidized PDIs catalyze the oxidation of substrate proteins by disulfide exchange resulting in disulfide formation. Subsequently, PDIs get re-oxidized by EROs, which in turn transfer electrons further to oxygen via an internal thiol-disulfide cascade (Aller and Meyer, 2013). Transferring two electrons to oxygen leads to incomplete reduction and thus the formation of hydrogen peroxide (H_2_O_2_), the fate of which is currently unclear. H_2_O_2_ may either leave the ER for detoxification in the cytosol or contribute to the oxidative power in the ER if it is reduced to water locally. Indeed, glutathione peroxidase-like 3 (GPXL3) has been identified in the ER and proposed to use H_2_O_2_ as an alternative oxidant for PDIs (Attacha et al., 2017; Meyer et al., 2021).

For correct folding of proteins, the formation of non-native disulfides needs to be avoided. Consequently, strict control and fine-tuning of the activity of the oxidizing machinery is essential. Typical PDIs have midpoint redox potentials in the range of -140 to -190 mV (Lundström and Holmgren, 1993; Matsusaki et al., 2016; Selles et al., 2017). Such comparatively oxidizing redox potentials would render PDIs thermodynamically highly active towards thiols in nascent peptides with the risk of generating erroneous disulfides. Reduced glutathione (GSH) counters the oxidation potential by maintaining a significant fraction of PDIs in the reduced state (Chakravarthi et al., 2006). Such thiol-disulfide exchange reactions typically operate close to their equilibrium, meaning that the *in vivo* potential of a thiol-disulfide redox couple can differ significantly from the midpoint potentials of the participating reactants. This implies that the resulting steady-state glutathione redox potential (*E*_GSH_) is a good approximation of the true redox potential of PDIs *in vivo*.

To complete the catalytic cycle, reduced PDIs will be re-oxidized by transferring electrons to EROs. In yeast ERO1 is the only ERO protein and mediates the re-oxidation of PDI and disulfide bond formation (Frand and Kaiser, 1998; Frand and Kaiser, 1999). A loss of ERO1 is lethal for yeast, proving it essential in the process of oxidative protein folding (Frand and Kaiser, 1998; Pollard et al., 1998). While most plant species, including Arabidopsis (*Arabidopsis thaliana*), contain two ERO homologs, few exceptions with three isoforms or only a single copy exist (Aller and Meyer, 2013; Fan et al., 2019; Meyer et al., 2019; Onda et al., 2009). In Arabidopsis, both single null mutants are viable but homozygous double mutants are lethal, indicating a degree of functional redundancy for both isoforms (Fan et al., 2019). In rice (*Oryza sativa*) that contains only a single-copy ERO1, RNAi-knockdown of *Os*ERO1 in the endosperm inhibited the formation of native disulfide bonds in proglutelins. Simultaneously the formation of proglutelin aggregates was promoted due to non-native intermolecular disulfide formation, indicating OsERO1 fulfills a crucial function in the maturation of storage proteins (Onda et al., 2009).

With oxygen as the ultimate electron acceptor of oxidative protein folding in the ER, insufficient oxygen availability under hypoxia is likely to impact heavily on this critical ER function. Hypoxic conditions arise in plants during waterlogging (Voesenek and Bailey-Serres, 2015), in distinct tissues like e.g. potato tubers (van Dongen and Licausi, 2015), or the hypoxic niches of meristematic tissues (Weits et al., 2019). In all cases, hypoxia compromises multiple cellular processes and mitochondrial respiration, and triggers the expression of specific sets of genes that allow plants to withstand hypoxic conditions (Schmidt et al., 2018; Wagner et al., 2019). Cellular hypoxia in plants has been assessed using different approaches ranging from Clark-type electrodes for oxygen concentration to transcriptional biosensors and probes built on knowledge about oxygen-dependent protein degradation (Licausi and Giuntoli, 2021). While hypoxia is known to cause increasing reduction of cytosolic NAD and an oxidative burst after re-oxygenation (Wagner et al., 2019; León et al., 2021), the consequences of hypoxia in the ER are unknown. Similarly, the effect of GSH on the ER redox homeostasis is poorly defined. Experimental treatment of plants with dithiothreitol (DTT) as a model inducer of thiol-based reductive stress causes an unfolded protein response (UPR) (Fuchs et al., 2022; Howell, 2013). While this likely occurs through direct interference disulfide formation or stability, the dynamics and degree of such reductive stress insults remains to be defined.

Redox-sensitive GFPs (roGFPs) enable dynamic monitoring of cellular redox conditions in live cells (Meyer and Dick, 2010; Schwarzländer et al., 2016). The probe variant roGFP2 with a midpoint redox potential of -280 mV has been shown to provide a readout of local *E*_GSH_ due to interaction with dithiol GRXs (Meyer et al., 2007; Trnka et al., 2020). The fusion of human Grx1 to roGFP2 has provided a sensor with high specificity for *E*_GSH_ and improved dynamic response (Gutscher et al., 2008). In contrast to the cytosol where roGFP2 is largely reduced, roGFP2 is almost fully oxidized when targeted to the ER lumen and thus cannot be used for dynamic redox imaging in the ER unless the probe is first reduced by DTT (Merksamer et al., 2008; Schwarzländer et al., 2008). Because pronounced responsiveness of roGFPs is limited to a range of about ±35 mV around their midpoint redox potentials, it can be concluded that the actual *E*_GSH_ in the ER lumen is at least 35 mV less negative than the midpoint potential of roGFP2 (Müller-Schüssele et al., 2021). For *in vivo* redox measurements in less reducing conditions, different variants of roGFP1 with midpoint redox potentials between -229 mV and -246 mV have been engineered (Lohman and Remington, 2008). Of these variants, roGFP1iL with a midpoint of -229 mV has been used for imaging the redox state in the ER of mammalian cells, albeit without a fused Grx domain for catalysis (van Lith et al., 2011). These measurements indicated a redox potential of -231 mV and unexplained light-dependent oxidation effects. Extraction of ER-targeted roGFP1iE (midpoint of -236 mV) and analysis of its redox state on protein gel blots has suggested an *E*_GSH_ in the ER of HeLa cells of -208 mV (Birk et al., 2013). With an alternative FRET-based probe design based on two fluorophores linked by a redox-sensitive domain with a midpoint potential of -143 mV, the *E*_GSH_ in the ER of Chinese hamster ovary cells was reported to be -118 mV (Kolossov et al., 2011). This extensive range of different redox potentials has left major ambiguity regarding steady-state levels of *E*_GSH_ in the ER. For plant cells, information on *E*_GSH_ and its dynamics in the ER has hitherto not been available.

To dissect the role of plant ERO proteins in setting the luminal redox conditions such that oxidative protein folding can occur, we generated Arabidopsis lines with diminished ERO activity. First, we challenged the redox systems of these mutants by addition of thiols or deprivation of oxygen as terminal electron acceptor. To dissect the observed phenotypes mechanistically we generated an optimized biosensor system for the ER by expressing Grx1- roGFP2iL in the lumen. Using a combination of reverse genetics and *in vivo* monitoring of redox changes with pharmacological approaches allowed us to dissect the role that EROs play in connecting different reducing and oxidizing inputs. The obtained data reveal that EROs act as major redox integrators to promote luminal redox homeostasis and set the base for further investigation of luminal redox responses during cellular stress.

## RESULTS

### Viable *ero1 ero2* double mutants are highly sensitive to reductive stress

To elucidate the functional importance of EROs in Arabidopsis, several T-DNA insertion lines for *ERO1* or *ERO2* were identified and characterized. For *ERO1* the mutant allele *ero1-3* (SALK_096805) was confirmed by semi-quantitative RT-PCR to be a null mutant (Supplemental Figure S1). ERO1 and ERO2 have a protein sequence identity of 67.8% and double null mutants are lethal (Fan et al., 2019). To generate mutants with diminished ERO activity, we searched specifically for non-null alleles of *ERO2*. The allele *ero2-3* carries a T-DNA in the 5’ UTR, 297 bps upstream of the start codon. Even though semi-quantitative transcript analysis indicated a slight decrease in transcript abundance (Supplemental Figure S1), this allele turned out to be too weak to justify further analysis. To overcome the lack of suitable double T-DNA insertion mutants with residual ERO activity, an artificial microRNA (amiRNA) targeting the sixth exon of *ERO2* was designed and constitutively expressed in both Arabidopsis WT and *ero1-3* (Supplemental Figure S2A, B). Quantitative real-time PCR analysis of *ERO1* transcripts confirmed that *ero1-3* is a null mutant. Expression of amiR*ERO2* in WT plants diminished *ERO1* transcript levels to 40–60% compared with the WT, indicating that amiR*ERO2* partially co-suppressed *ERO1* (Supplemental Figure S2C). *ERO2* expression in *ero1-3* was not significantly different compared to WT (*P* = 0.1994, ANOVA with Dunnett’s multiple comparison test) indicating the lack of compensatory effects. Expression of amiR*ERO2* in WT plants caused a decrease of *ERO2* transcript levels down to 11–16% compared to WT, confirming the functional efficiency of the knockdown construct. Surprisingly, the decrease in *ERO2* transcripts was less pronounced in the *ero1-3* background, possibly due to compensatory gene regulation in response to the presumably severely diminished ERO activity. The respective double mutants, denominated as *ero1 ero2,* had transcript amounts for *ERO2* of 19–38% compared to the WT but did not show a pronounced phenotype under normal growth conditions (Supplemental Figure S2E-H). For further phenotypic analysis, the lines amiR*ERO2*#5 and *ero1 ero2*#5 were selected based on the lowest transcript abundance for *ERO2*.

Expression of *ERO1* and *ERO2* genes has been reported to be upregulated in plants exposed to pharmacological treatments that induce the unfolded-protein response (UPR), emphasizing the role of the respective proteins in protein-folding capacity (Fan et al., 2019). In line with findings from Fan et al., we found that upon incubation with 2 mM dithiothreitol (DTT) or 5 µg/mL tunicamycin (TM) *ERO1* transcript abundance increases several-fold. In our hands, however, induction of *ERO2* was observed neither with TM nor with DTT. This result might be linked to the UPR element, which is present only in the *ERO1* promoter sequence (Supplemental Figure S3).

Single null mutants for *ERO1* and *ERO2* are sensitive to reductive stress imposed by high DTT concentrations (Fan et al., 2019). To test the sensitivity of the *ero1 ero2* double mutant generated in this work, we germinated seeds of WT, *ero1-3*, amiR*ERO2* and the *ero1 ero2* on MS plates for 5 days and then transferred the seedlings to MS plates supplemented with 0–2000 µM DTT (Figure 1; Supplemental Figure S4). Root growth was already partially inhibited by 450 µM DTT in all tested genotypes and became more severely inhibited with higher DTT concentrations. While *ero1-3* seedlings became more sensitive to DTT than WT seedlings only at 2 mM DTT, *ero1 ero2* displayed a pronounced sensitivity already on 450 µM DTT (Figure 1C) and even 200 µM DTT in a side-by-side comparison with WT seedlings (Supplemental Figure S4). Interestingly, primary root growth of amiR*ERO2* seedlings was not more sensitive to DTT than the WT even at 2 mM DTT (Figure 1C). Taken together, these results support the notion that both ERO isoforms have partially redundant functions in redox-based processes in the ER lumen of Arabidopsis and loss of one isoform can be partially compensated by the other isoform.

**Figure 1.**
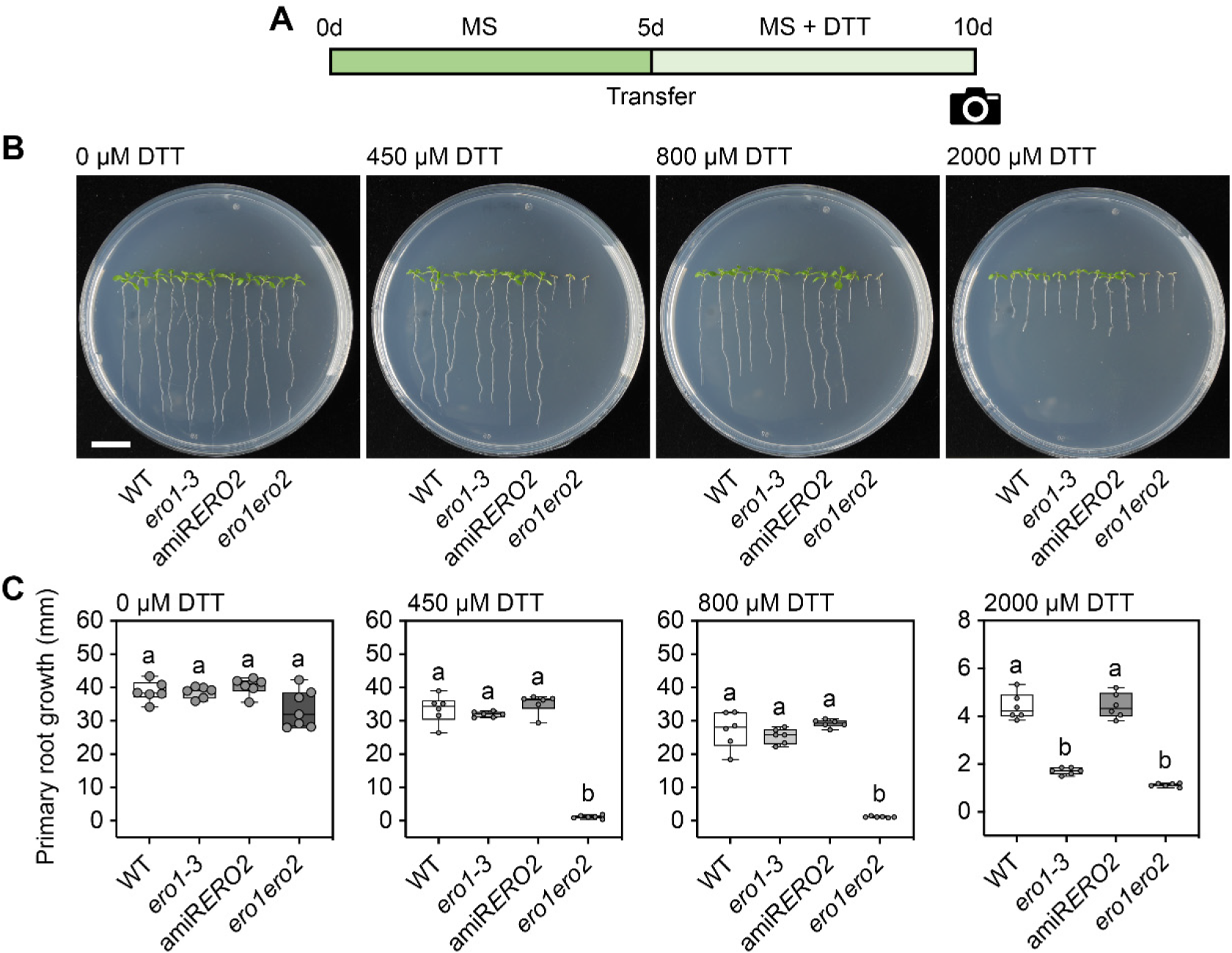
*ero1 ero2* seedlings are hypersensitive to reductive stress. A, Scheme indicating the reductive stress treatment performed on seedlings. Seedlings were grown for 5 days on MS agar plates before transferred to MS agar plates supplemented with different concentrations of DTT. After growth for 5 additional days, seedlings were documented and primary root growth after transfer assessed. B, Representative images of WT, *ero1-3*, amiR*ERO2* and *ero1 ero2* seedlings after exposure to the indicated DTT concentrations. Bar = 1 cm. C, Primary root growth of WT, *ero1-3*, amiR*ERO2* and *ero1 ero2* seedlings after transfer to the indicated DTT concentrations, *n* = 6-33. Box = interquartile range between the lower and upper quartiles, center line = median, whiskers = min and max values. Statistical analyses were performed using ANOVA with Fisher’s LSD test. Different letters indicate statistically different groups (*P*<0.05). Severe DTT sensitivity of *ero1 ero2* mutants is also supported by Supplemental Figure S4.

### ERO activity is essential for hypoxia tolerance

ERO activity requires molecular oxygen as the ultimate acceptor for electrons released during disulfide formation on nascent peptides imported into the ER. It has been hypothesized that ERO activity might be severely compromised under hypoxia, which may lead to accumulation of unfolded proteins and induction of UPR (Schmidt et al., 2018; Meyer et al., 2019). To test whether a bottleneck in the ER redox systems by diminished ERO capacity results in higher sensitivity to oxygen limitation, WT and *ero1 ero2* plants were exposed to different hypoxia regimes (Figure 2). When seedlings grown on MS plates were deprived of oxygen for 8.5 h followed by a 3-day recovery phase, seedlings were impaired in growth and showed chlorotic leaves (Figure 2A). Categorization of each seedling into one of three injury classes and quantitative evaluation showed that *ero1 ero2* mutants had a lower survival score than WT seedlings (*P* = 0.0007, unpaired t-test) (Figure 2A–C). Susceptibility to hypoxia was also observed in soil-grown plants at rosette stage after flooding for 4 or 5 days. In general, submergence decreased vegetative growth of the rosette and triggered cell death for both genotypes, but *ero1 ero2* was consistently more severely affected than WT plants after 4 d (WT = 0.46 **±** 0.81 g; *ero1 ero2* = 0.31 **±** 0.14 g) and 5 d of submergence (WT = 0.38 **±** 0.06 g; *ero1 ero2* = 0.21 **±** 0.08 g) (Figure 2D, E). Diminished hypoxia tolerance of *ero1 ero2* seedlings was also observed in primary roots of 5-day-old seedlings after intermittent exposure to ∼0.1% O_2_ for three hours in an anaerobic chamber. After a 48-h recovery period under normoxia, 37.8% of WT seedlings but only 16.7% of *ero1 ero2* had resumed root growth (*P* = 0.0051, unpaired t-test) (Figure 2F, G).

**Figure 2.**
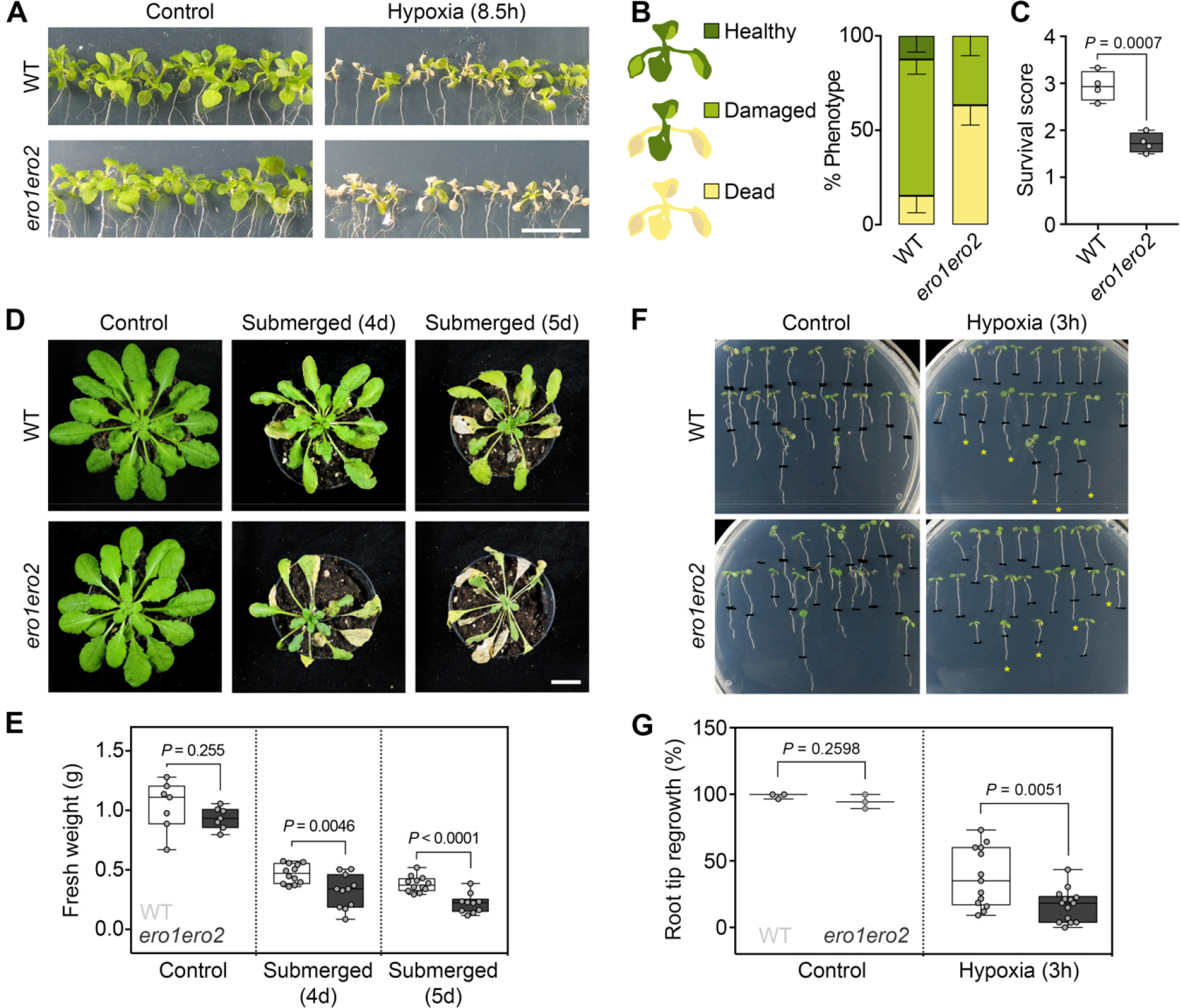
ERO activity is required to tolerate hypoxic conditions. A–C, Impact of hypoxia on the survival of WT and *ero1 ero2* seedlings. 11-day old seedlings on MS plates were exposed to < 0.3% O_2_ for 8.5 h and recovered for 3 days under normoxia and long-day conditions (A). According to the phenotype severity after recovery, seedlings were classified into three categories (healthy, damaged or dead) (B). Based on the number of seedlings in each category, a survival score was calculated, *n* = 4 (C). D–E, 6-week-old WT and *ero1 ero2* plants grown in soil were submerged in water for 4 or 5 days in the dark and recovered for 6 days under normoxia and a short-day illumination regime (D). After treatment, the fresh weight from rosettes was measured, *n* = 7–12 (E). F–G, Impact of hypoxia on primary root growth. Five- day-old WT and *ero1 ero2* seedlings on MS plates were exposed to ∼ 0.1% O_2_ for 3 h in an anaerobic chamber and subsequently allowed to recover in normoxia for 48 h. Seedlings that were able to continue primary root growth after hypoxia (marked with * in panel F) were counted and expressed relative to the total number of seedlings (G) *n* = 3–14. Box = interquartile range between the lower and upper quartiles, center line = median, whiskers = min and max values. Statistical analyses were performed using unpaired t-test.

### ERO1 and ERO2 are type II ER membrane proteins

While different bioinformatic algorithms consistently predict ERO1 to be targeted to the secretory pathway, predictions for ERO2 are less uniform and include plastids and the nucleus besides the ER. In contrast to their soluble mammalian counterparts and yeast Ero1p, which is a type I protein with a C-terminal transmembrane domain (TMD), both Arabidopsis EROs are predicted to be type II proteins with a TMD close to their N-termini (ARAMEMNON database, http://aramemnon.uni-koeln.de/; (Schwacke et al., 2003). To experimentally verify localization and topology, we performed a redox-based topology assay (ReTA, (Brach et al., 2009) by expressing both ERO1 and ERO2 tagged at either their N- or their C-termini with roGFP2 in leaves of *Nicotiana benthamiana*. Given the steep gradient in *E*_GSH_ across the ER membrane, roGFP2 is expected to be fully reduced on the cytosolic side and fully oxidized when facing the ER lumen, respectively. In consequence, a binary readout is expected for N- and C-terminal roGFP2 fusions of a single-spanning membrane protein. Expression of N- and C-terminal fusions of both EROs with roGFP2 resulted in a network-like pattern including a nuclear ring that co-localized with the ER-marker, AtWAK2_TP_-mCherry-HDEL (Figure 3). This suggests that both proteins were targeted to the secretory pathway and that even tagging their N-termini with roGFP2 did not mask the respective targeting signals.

**Figure 3.**
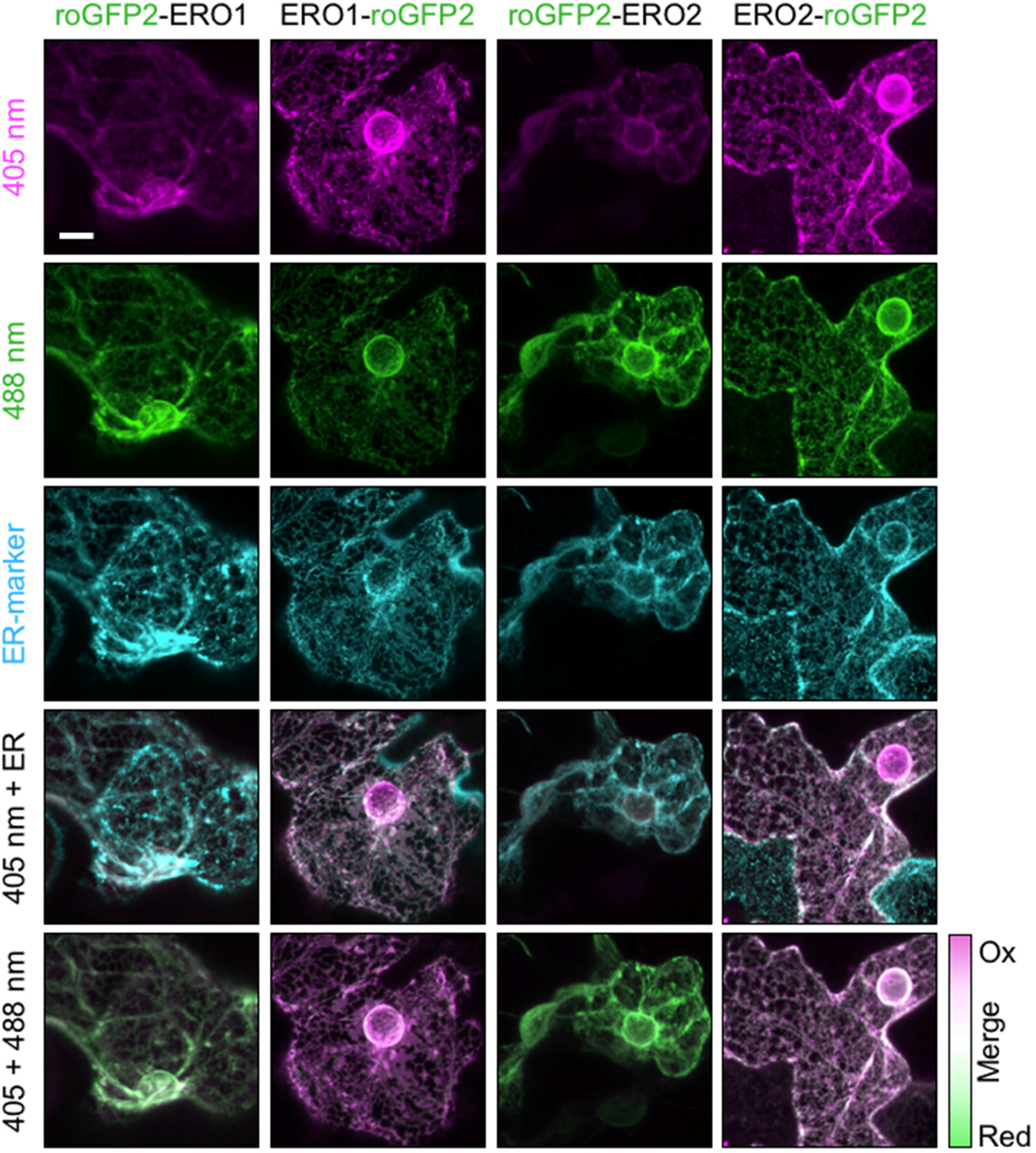
Analysis of ERO1 and ERO2 protein orientation in the ER membrane. Confocal images of the fluorescence signal from *Nicotiana benthamiana* leaf epidermal cells transiently co-transformed with fusions of roGFP2 to either the N- or the C- termini of ERO1 or ERO2, and the ER-marker AtWAK2_TP_-mCherry-HDEL (Nelson et al., 2007). roGFP2 fluorescence was collected at 505–530 nm after excitation with either 405 nm or 488 nm. The ER-marker was excited at 543 nm and the fluorescence collected at 590–630 nm. Scale bar = 10 µm.

Due to the large dynamic range of roGFP2, a simple merge of the false color-coded images collected with 405 nm and 488 nm excitation, respectively, was sufficient to gain a binary readout for the orientation of EROs in the membrane. For both EROs, the N-termini are facing the cytosol while the C-termini are oriented towards the ER lumen. This unambiguously identifies both EROs as type II membrane proteins (Figure 3; Supplemental Figure S5).

### Sensing the glutathione redox potential in the ER lumen with Grx1-roGFP2iL *in vivo*

To maintain homeostatic redox conditions in the lumen the reducing power generated by the import of nascent peptides into the ER must be matched stoichiometrically by the oxidizing power of EROs as disulfide-generating proteins. We hypothesized that any disruptions to rates in electron flow though the redox systems involved will give rise to imbalance, i.e. dynamic redox changes. To test this hypothesis, it is necessary to visualize dynamic redox changes specifically in the ER lumen. Based on the assumption that GSH in the ER lumen contributes to the local redox balance and that an equilibrium between *E*_GSH_ and the redox potential of putative protein disulfides is established, we thus explored possibilities for developing an appropriate biosensor for the luminal *E*_GSH_. roGFP2 is completely oxidized in the ER and thus cannot sense dynamic changes that may occur in stress situations. We thus turned to roGFP2iL, which has a less negative midpoint potential of -240 mV compared to -280 mV for roGFP2 (Aller et al., 2013). roGFP2iL was fused to Grx1 to ensure rapid equilibration with the local *E*_GSH_ (Gutscher et al., 2008) and was equipped with the C-terminal ER retrieval signal HDEL. This sensor construct was fused with the targeting peptide (TP) from Arabidopsis chitinase and then expressed from a UBQ10 promoter (Supplemental Figure S6A). Transient expression of this construct in *N. benthamiana* leaves in combination with the ER-marker AtWAK2_TP_- mCherry-HDEL confirmed that the sensor was targeted and retained in the ER as expected (Supplemental Figure S6B).

Stable expression of Grx1-roGFP2iL-HDEL in Arabidopsis resulted in a typical reticulate ER network labeling including ER bodies that can be recognized as intensely fluorescent elongated structures (Figure 4A; Supplemental Figure S7). When expressed and imaged in WT, *ero1-3* or amiR*ERO2* seedlings, the ratio analysis resulted in very similar ratio values with a slightly lower ratio of *ero1-3* compared to amiR*ERO2* (Supplemental Figure S7). A more pronounced drop in the fluorescence ratio, indicative of a reductive shift, was found in the ER of *ero1 ero2* (Figure 4; Supplemental Figure S7). For calibration and evaluation of the sensor responsiveness, seedlings of WT and *ero1 ero2* were immersed in either 10 mM DTT, 100 mM H_2_O_2_, or in water as control (Figure 4A–D). In both genetic backgrounds, the sensor was fully responsive with δ between 2.0 and 2.2, which is similar to the δ obtained in tobacco cells (Supplemental Figure S6C, D). Based on the calibration values for fully reduced and fully oxidized sensor the ratio values were converted to the degree of sensor oxidation. While *OxD_Grx1-roGFP2iL_* in WT was 46.5% the *OxD* in the ER of *ero1 ero2* was only 25.3% (Figure 4E). Direct comparison of these *OxD* values with the titration curve for Grx1-roGFP2iL calculated from the Nernst-equation suggests *E_GSH_* values of -241 mV for the ER of WT plants and -253 mV for *ero1 ero2* (Figure 4F). Considering that the midpoint potential of Grx1-roGFP2iL matches WT *E*_GSH_ the sensor is exquisitely well-suited for dynamic redox measurements in the ER being able to reliably capture redox changes towards oxidation and reduction.

**Figure 4.**
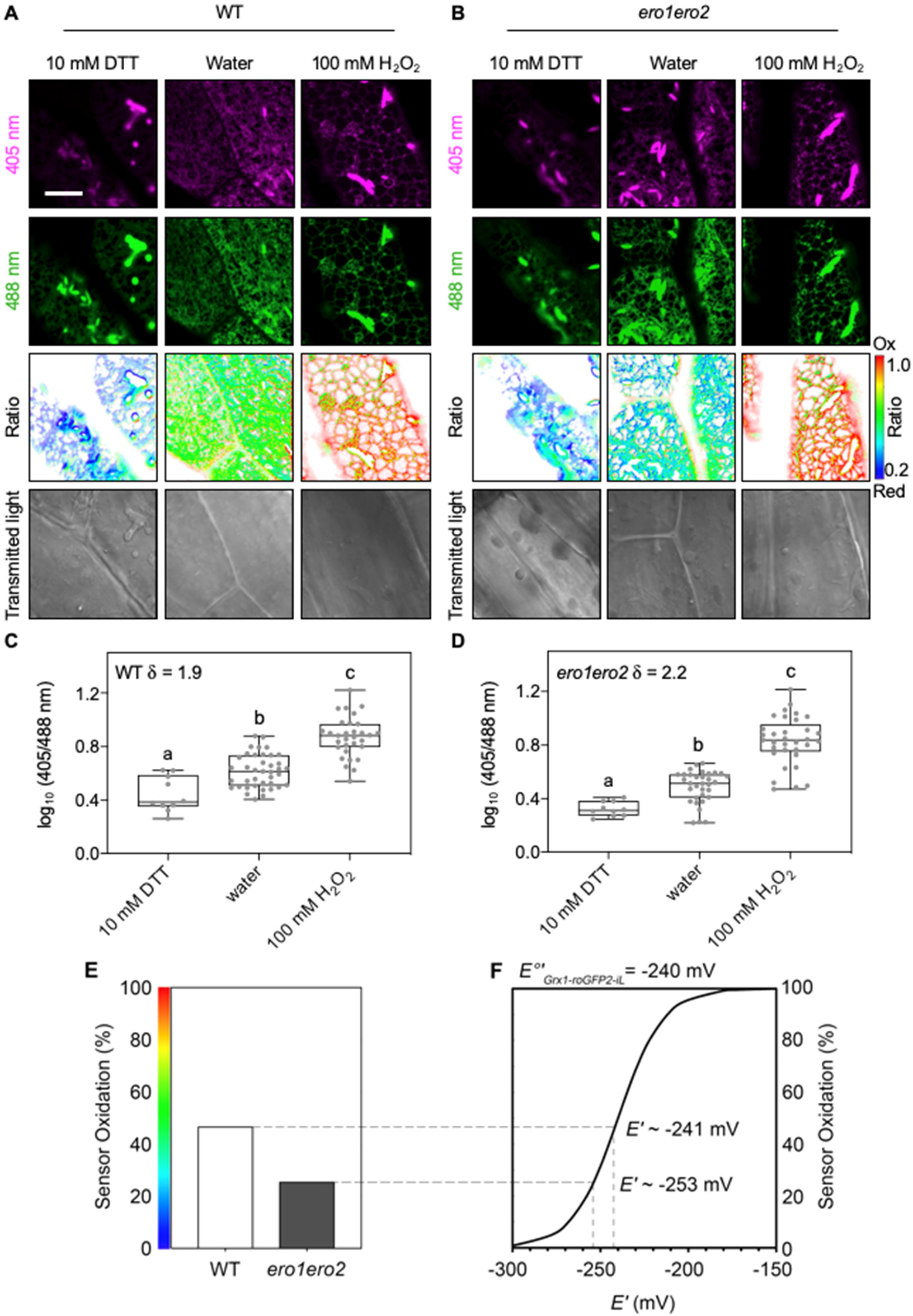
*In vivo* monitoring of the glutathione redox potential reveals less oxidizing conditions in the ER of *ero1 ero2*. A-B, Confocal images of hypocotyl cells of five-day old Arabidopsis WT (A) and *ero1 ero2* (B) seedlings stably expressing the Grx1-roGFP2iL-HDEL. roGFP2iL fluorescence was collected at 505–530 nm after excitation with either 405 nm or 488 nm. Ratio images were calculated as the 405 nm/488 nm fluorescence. To fully reduce or oxidize the sensor, seedlings were immersed in 10 mM DTT or 100 mM H_2_O_2_, respectively. Control samples were immersed in deionized water as control. False colors indicate the fluorescence ratio on a scale from blue (reduced) to red (oxidized). Scale bar = 10 µm. C-D, Fluorescence ratios for WT (C) and *ero 1ero2* (D), *n* = 10–39. δ = dynamic range of the sensor. Box = interquartile range between the lower and upper quartiles, center line = median, whiskers = min and max values. Statistical analyses were performed using ANOVA with Fisher’s LSD test. Different letters indicate statistically different groups (*P*<0.05). E, Sensor oxidation *OxD* for Grx1-roGFP2iL-HDEL in hypocotyl cells from WT and *ero1 ero2* calculated from the mean fluorescence data shown in panels C and D. F, Redox titration curve for roGFP2iL calculated from the Nernst-equation with *E_0_’_Grx-roGFP2iL_* = -240 mV. Interpolation of the *OxD* in panel E within the titration curve suggests that *E*_GSH_ in the ER of WT lines is about -241 mV, and -253 mV for *ero1 ero2*. Redox measurement in further *ero* mutants is shown in Supplemental Figure S7.

### Grx1-roGFP2iL imaging reveals ERO-dependent *E*_GSH_ dynamics in the ER lumen after a reductive challenge

Both, stress situations and physiological situations with intense secretion may affect the redox balance in the ER and demand dynamic readjustments. To test for the ability of Grx-roGFP2iL to sense dynamic changes in the luminal *E*_GSH_ and to further explore the role of ERO activity in redox readjustments, we evaluated the capacity of the ER to restore its redox homeostasis after a pulse of reductive stress. WT and *ero1 ero2* seedlings expressing Grx1-roGFP2iL-HDEL were mounted in a perfusion chamber on the microscope stage and exposed for approximately one minute to 5, 50 or 500 µM DTT followed by perfusion with imaging buffer (Figure 5A, B). Dynamic changes in sensor fluorescence were followed for the root elongation zone where direct comparison of data for WT and *ero1 ero2* mutants highlighted lower 405 nm/488 nm ratios in the *ero1 ero2* mutant (Figure 5C). This finding is consistent with a lower ratio observed in hypocotyl cells of *ero1 ero2* seedlings compared to the wild-type (Figure 4). After initial recording of steady-state values, perfusion of roots with DTT caused a dose-dependent decrease of fluorescence ratios for both genotypes with a comparable reduction rate. The reduction rate was calculated as the linear slope of the graph (ratio change per time) in the initial phase of DTT perfusion (Figure 5D). Maximum reduction depicted as the lowest recorded fluorescence ratio was the same for 50 and 500 µM DTT shortly after start of the DTT washout. Because the decrease in fluorescence ratios was transient without reaching a plateau, the lowest ratio values are unlikely to represent the fully reduced state of the sensor (Figure 5A, B). Immediately after effective washout of DTT from the perfusion chamber, i.e. about 1 min after switching from DTT to imaging buffer, fluorescence ratio values in all samples started to recover to reach values similar to values before DTT perfusion. The re-oxidation rate calculated as the linear slope after DTT washout was lower for *ero1 ero2* than for WT (Figure 5C, E). Together these results show that Grx1-roGFP2iL is suitable for dynamic measurements of *E*_GSH_ in the ER. Furthermore, the data suggest that the luminal *E*_GSH_ is directly dependent on ERO activity and that ERO activity contributes to redox homeostasis in the ER lumen both to maintain a defined steady state and to efficiently re-establish that steady-state after a reductive challenge.

**Figure 5.**
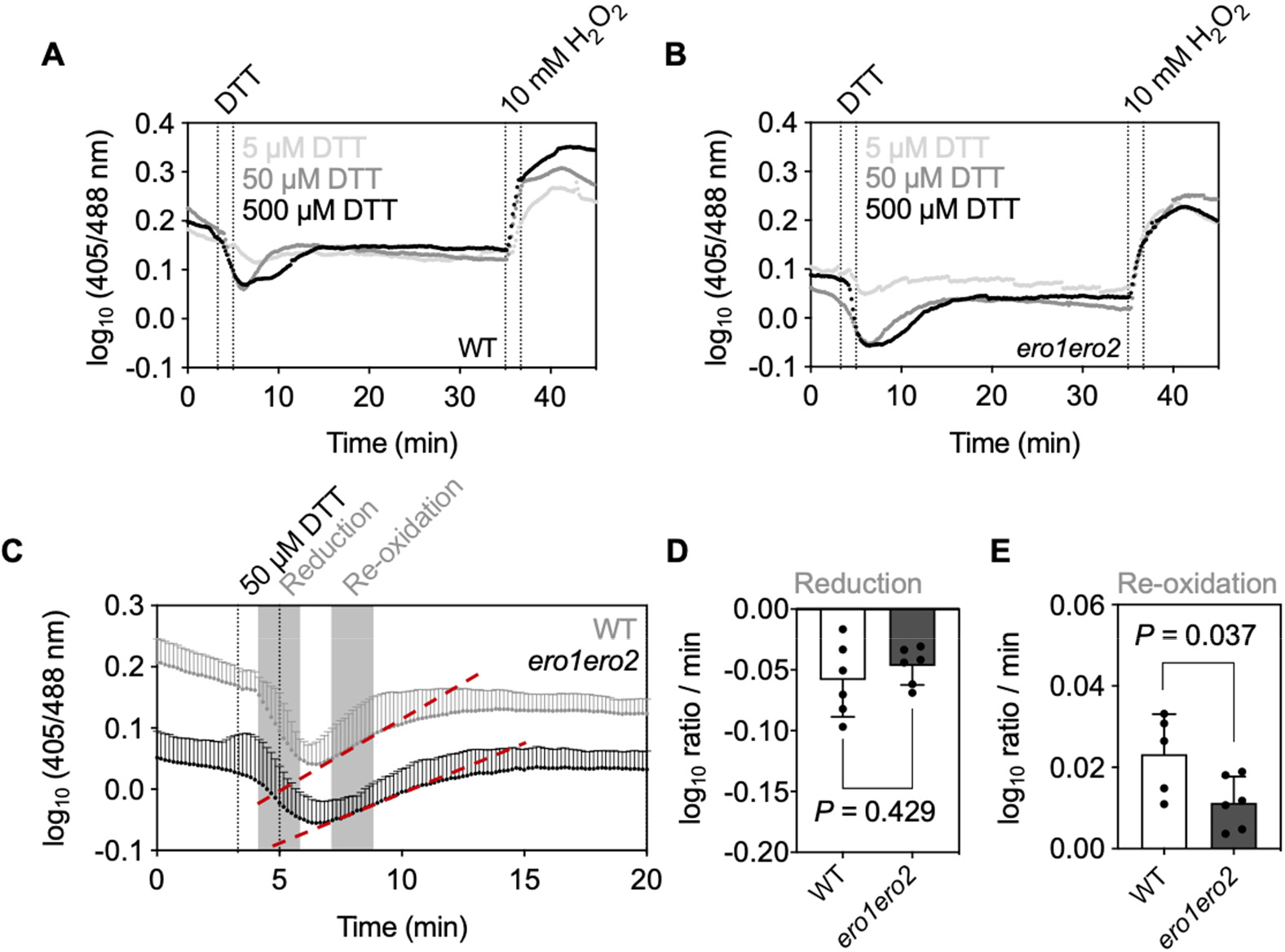
ERO activity is required to reestablish the ER redox homeostasis after DTT- induced reductive stress. A-B, Four-day-old Arabidopsis WT (A) and *ero1 ero2* (B) seedlings stably expressing Grx1-roGFP2iL-HDEL were mounted in a perfusion chamber on the stage of a confocal microscope. Cells in the root elongation zone were imaged while the samples were continuously perfused with imaging buffer, DTT (5–500 µM) or 10 mM H_2_O_2_. roGFP2iL fluorescence was collected at 505–530 nm after excitation with either 405 nm or 488 nm. Log_10_ values of the 405 nm / 488 nm fluorescence ratio were calculated and plotted for all indicated treatments. The presented data indicate the mean log_10_ ratio with *n* = 3. C, Direct comparison of mean log_10_ ratios + SD of WT and *ero1 ero2* seedlings perfused with 50 µM DTT. Dotted lines indicate the time window in which seedlings were perfused with DTT. Pronounced sensor reduction occurred only after a delay of ∼1 min, which is due to the dead volume of tubing and the perfusion chamber. Grey areas indicate the time windows in which the speed of reduction and oxidation was measured. For the re-oxidation, this is depicted by dashed red lines drawn as tangents to the ratio plot in this response phase. D–E, Rates for sensor reduction after DTT perfusion (D) and re-oxidation after washout of DTT (E) from root tissues calculated from the time course data in panel C. Bars indicate the mean rates - SD (D) or + SD (E).

### ERO activity influences luminal *E*_GSH_ dynamics in response to hypoxia

With O_2_ being the final electron acceptor in ERO-mediated oxidative protein folding in the ER, hypoxia-induced restriction of oxidizing power may be expected to affect the luminal *E*_GSH_. Based on the initial observation that plants deficient in ERO activity are more susceptible to low O_2_ conditions than WT plants (Figure 2), we assessed whether and to what extent hypoxia affects ER redox homeostasis and how ERO activity contributes to establishing new redox equilibria under such conditions.

To establish hypoxic atmospheric conditions and to enable simultaneous readouts for multiple replicates, we used a plate reader setup with a build-in atmospheric control option and used leaf discs immersed in imaging buffer on 96-well plates as a technical advancement of an approach that we had optimized previously (Wagner et al., 2019). For initial evaluation of the effects of severe hypoxia on *E*_GSH_ in the ER, we exposed leaf disks of WT plants expressing Grx1- roGFP2iL-HDEL to an atmosphere with only 0.1% O_2_ for 1, 3.5 or 6.5 h and followed the sensor response by monitoring the fluorescence excited at 400 nm and 480 nm, respectively, as well as the resulting 400 nm / 480 nm fluorescence ratio (Supplemental Figure S8). Immediately after O_2_ depletion, the sensor responded with decreased fluorescence excited at 400 nm and a concomitant increase in fluorescence excited at 480 nm. The respective log_10_(400 nm/480-nm ratio) values indicate successive reduction of the probe. This reduction, however, lasted only for about 45 min and was followed by a gradual re-oxidation even when the hypoxic phase was extended to 3.5 and 6.5 hours. For the 3.5-h hypoxic phase, increasing fluorescence excited at 400 nm and decreasing signal after excitation at 480 nm indicated reliable recording of pronounced sensor oxidation upon re-oxygenation (Supplemental Figure S8C. D), which led us to do all further experiments with a 3.5 h hypoxic treatment.

Exposure of *ero1-3* and amiR*ERO2* to hypoxia led to ratio traces for luminal Grx1-roGFP2iL- HDEL very similar to the WT. In all cases, the initial hypoxia-induced reduction was followed by a gradual ratio increase during the hypoxic phase and a transient oxidative upon re- oxygenation (Figure 6A). In the ER of *ero1 ero2* mutants, however, the initial ratio values at the start of the experiments were already lower than in the other lines (Figure 6A, B). This is consistent with the measurements of steady-state *E*_GSH_ values by confocal microscopy (Figure 4). Despite the reductive shift at steady state, the sensor in *ero1 ero2* responded to hypoxia with the initial reduction very similar to all other lines (Figure 6A, B). The subsequent gradual ratio increase during the hypoxia phase did not occur. Re-oxygenation in this case also caused a ratio increase, but gradually approaching the original steady-state values from reduced values without a pronounced transient peak immediately after re-oxygenation (Figure 6A, C).

**Figure 6.**
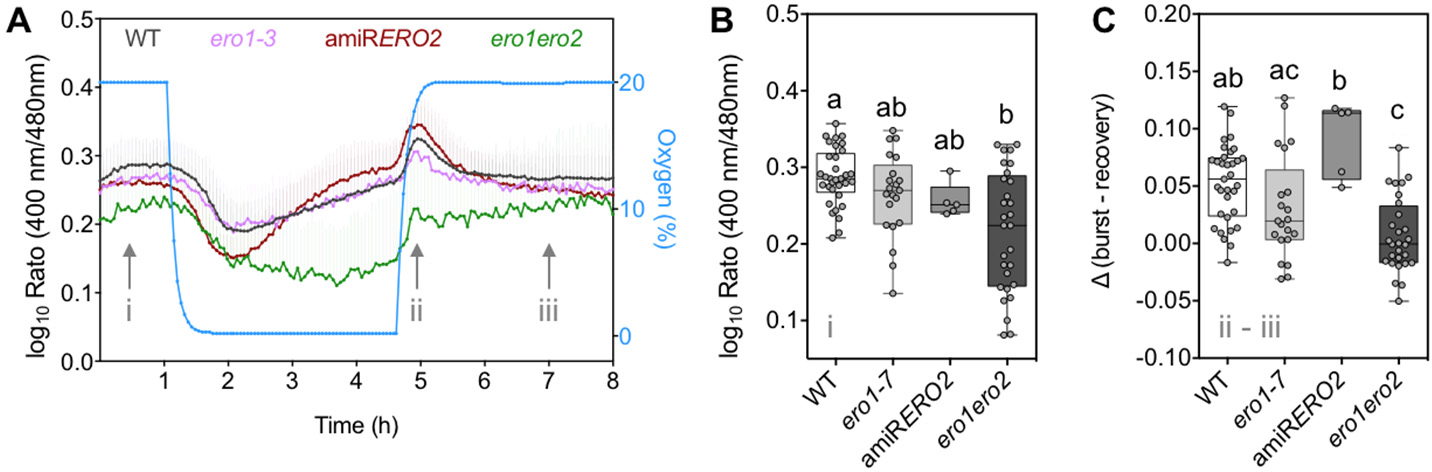
*ERO1* and *ERO2* are required to maintain ER redox homeostasis under low oxygen conditions. A, Leaf disks from four-week-old WT, *ero1-3*, amiR*ERO2* and *ero1 ero2* plants expressing the sensor Grx1-roGFP2iL-HDEL were placed in 96-well plates, immersed in 200 µL imaging buffer. Plates with samples and non-fluorescent controls were place in a plate reader equipped with an atmospheric control unit that enabled fast changes of atmospheric conditions. roGFP2iL fluorescence was continuously collected at 520 ± 10 nm after excitation at 400 ± 5 nm and 480 ± 5 nm. Sensor fluorescence was measured for 1 h at 20% O_2_ (normoxia), before O_2_ was decreased to 0.1% (hypoxia) for 3.5 h. Subsequently, normoxic conditions were reestablished and fluorescence monitored for further 3.5 h. Traces shows mean values for log_10_- transformed 400 nm / 480 nm fluorescence ratios + SD, *n* = 5-33. B, Steady state fluorescence ratios at the start of the experiment (i). C, Difference in log_10_-transformed fluorescence ratios at the peak of the re-oxygenation (ii) burst and its posterior recovery (iii). Box = interquartile range between the lower and upper quartiles, center line = median, whiskers = min and max values. Statistical analyses were performed using ANOVA with Tukey’s multiple comparison test. Different letters indicate statistically different groups (*P*<0.05).

### The luminal *E*_GSH_ integrates different reducing and oxidizing inputs

We next attempted to dissect the different reducing and oxidizing inputs that shape the characteristic redox dynamics of the ER lumen during the hypoxia regime. Based on the observation of the initial reduction after onset of hypoxia, we hypothesized that the net increase in reductive pressure results from a shift in the balance between continuous import of nascent peptides with reduced thiols and the oxidative power generated by EROs on the other hand. At the sharp drop in O_2_ supply, ERO activity is expected to stop while the reducing input from nascent peptides may persist for longer. To test the hypothesis, leaf disks where pre-incubated for 18 h with 70 µM cycloheximide (CHX) as an inhibitor for protein synthesis. Treatment of leaf discs from WT plants with CHX to stop *de novo* protein synthesis and intake of protein thiols into the lumen caused more oxidized steady-state values in the lumen, which is consistent with the concept that the steady-state ratio values of Grx1-roGFP2iL-HDEL reflects the balance between oxidative and reductive processes (Figure 7A). This increase in the steady-state ratio was absent in *ero1 ero2* (Figure 7B), consistent with the idea that EROs provided the oxidation power responsible for the oxidation in the CHX-treated WT tissue. In both genetic backgrounds, WT and *ero1 ero2*, CHX abolished the transient reduction during the initial hypoxia phase and the subsequent gradual ratio increase that is normally seen in the WT (Figure 7A, B). These observations taken together suggest that the initial reductive drop under hypoxia is caused by continued import of nascent peptides into the ER. While re-oxygenation in wild-type leaf discs abolished the pronounced sensor oxidation after re-oxygenation, the fluorescence ratio values still increased with a delay and subsequently returned to steady-state values more slowly than in leaf discs not treated with CHX (Figure 7A).

**Figure 7.**
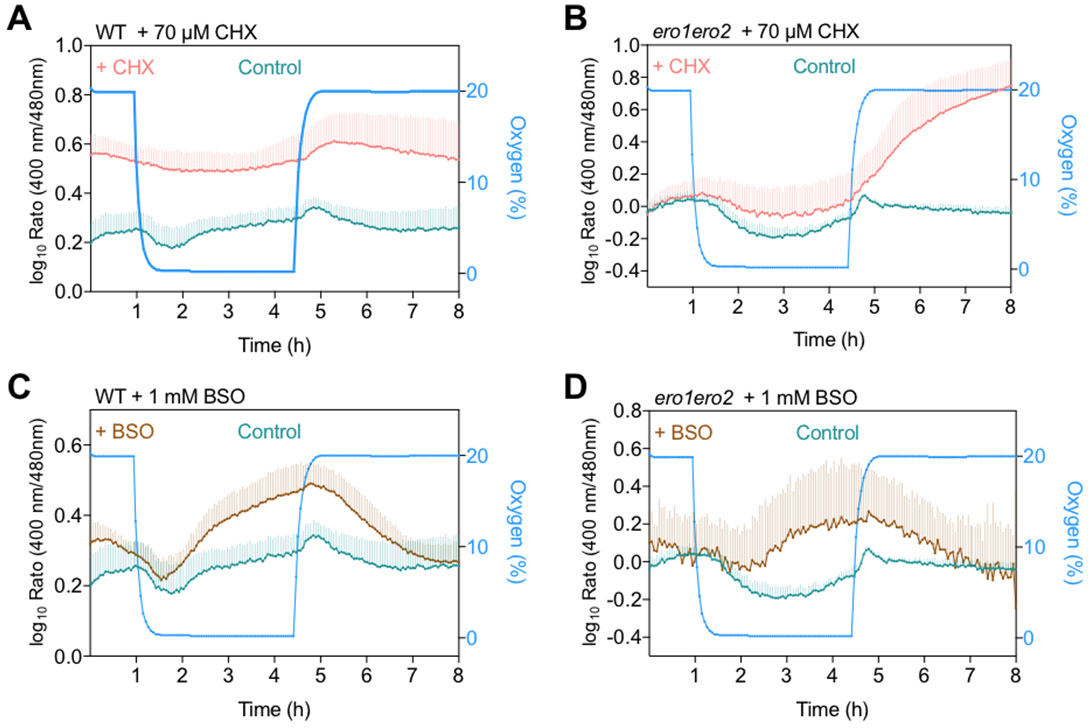
*E*_GSH_ in the ER reflects the equilibration of reducing and oxidizing inputs. A-D, Effect of cycloheximide (CHX, 70 µM) (A, B) and buthionine sulfoximine (BSO, 1 mM) (C, D) on hypoxia-induced redox dynamics in the ER of WT and *ero1 ero2* leaf disks. In all cases, leaf disks immersed in imaging buffer were used as controls. Controls in panels C and D are the same as in panels A and B, respectively, because data for CHX and BSO were collected on the same plate. In CHX-treated *ero1 ero2* mutants, the re-oxygenation led to a gradual log_10_ ratio change from about 0 to 0.75 (B). This change, however, would be equivalent to a δ of about 5.6 and thus much larger than what would be expected for a redox response of roGFP2iL. Hence, the observed response in this case very likely indicates some unknown underlying factor that compromises the sensor response. The normalized fluorescence for individual channels can be found in Supplemental Figure S9.

Besides import of nascent peptides, reductive power in the ER may also originate from continuous import of GSH as the second major cellular pool of thiol-based reductant. To test this possibility, leaf disks of WT and *ero1 ero2* were pre-incubated in 1 mM buthionine sulfoximine (BSO) as a specific inhibitor of glutamate-cysteine ligase (GSH1), which catalyzes the first step in GSH biosynthesis (Meyer and Fricker, 2002). Due to continuous turnover of GSH, the induced lack of resupply results in gradually decreasing GSH levels. Steady-state ratio values after 18 hours pre-incubation with BSO were already slightly higher than in control samples (Figure 7C, D). The gradual ratio increase during the second phase of the hypoxia treatment was far more pronounced in BSO-incubated leaf discs than in control samples in both WT and *ero1 ero2* mutants (Figure 7C, D). After re-oxygenation, the ratios gradually decreased and returned to values similar to the starting values (Figure 7C, D). This suggests that after re- oxygenation import of nascent polypeptides into the ER resumes, dominating the redox landscape in the lumen.

## DISCUSSION

### Diminished activity of ERO renders plants highly sensitive to reductive stress and hypoxia

Functional redundancy between proteins of the same family is a common feature in plants, and that can compromise straightforward functional analysis by reverse genetics. In the case of Arabidopsis ERO, where two gene loci show conservation with the mammalian and yeast EROs, individual null mutants of either gene were isolated and described show hypersensitivity to 2 mM DTT (Fan et al., 2019). While neither of the null mutants showed a distinct phenotype, the genetic combination of both turned out to be lethal. The *ero1 ero2* double mutant generated here circumvents this problem by abolishing *ERO1* transcript and diminishing the amount of *ERO2* transcript. Even though the *ero1 ero2* plants show no obvious phenotype under non-stress lab conditions, the seedlings were hypersensitive to DTT exposure, which acts as a potent thiol-based reductant that readily enters the cell and the ER (Figure 1). In biochemical assays for ERO proteins, DTT is frequently used as a substrate (Beal et al., 2019; Fan et al., 2019). The DTT-sensitivity suggests that *in vivo* EROs are required to directly counter the reductive impact by oxidizing DTT to avoid deleterious effects of DTT on existing disulfides on proteins. The hypersensitivity to reductive stress also provides direct evidence for the presence of both isoforms in Arabidopsis roots and a degree of overlap in their biochemical function. Despite containing only one *ERO* gene, knockdown mutants of rice did not show any other phenotype beyond aggregated proglutelins, which could theoretically indicate a specific requirement for ERO for proglutelin folding. Considering our results as well as previous work in yeast and mammalian systems on ERO function, it seems more plausible, however, that ERO is required for oxidation of multiple proteins entering the secretory pathways and that proglutelin aggregation is caused by an overload of the remaining oxidative capacity of the ER. The correct folding of different target proteins may have distinct thresholds for minimal ERO activity. A different threshold could also be defined by a low-efficient alternative oxidation system similar to alternative oxidation pathways described in mice with peroxiredoxin IV (Tavender et al., 2010; Zito et al., 2010), glutathione peroxidase (Nguyen et al., 2011; Wang et al., 2014), and vitamin K epoxide reductase (Schulman et al., 2010). While such systems may enable limited disulfide formation without ERO, they cannot explain the observed hypersensitivity of *ero1 ero2* mutants to hypoxia. In yeast, the fumarate reductase Osm1 that is dual localized in the intermembrane space of mitochondria and the ER, can transfer electrons from FAD to fumarate (Neal et al., 2017; Kim et al., 2018). In the absence of oxygen, Osm1 may accept electrons from the Ero1 FAD, allowing *de novo* disulfide bond formation under anaerobiosis (Kim et al., 2018). Through this system, electrons from luminal disulfide formation may be redirected to mitochondria. Although no such alternative oxidation system has been identified yet in Arabidopsis, our observation of increased hypoxia sensitivity of *ero1 ero2* may be a first hint at the existence of an alternative, albeit less efficient, acceptor for electrons from EROs and ultimate transfer to acceptors in other compartments. Recently, we showed that mitochondrial respiration can act as a backup system for thiol-based reductants that enter the ER from the outside (Fuchs et al., 2022). Whether a surplus of thiol-derived electrons accumulating in the ER under hypoxia can be redirected in a similar way awaits further analysis in the future. Irrespective of the existence of a low-efficient backup, it is surprising how quickly hypoxic conditions become deleterious and largely abolish any recovery after only a few hours (Figure 2). Plant cells exposed to hypoxia quickly run into an energy crisis due to the lack of the final electron acceptor for mitochondrial respiration reflected by a decrease in ATP levels and concomitant increase in the NADH/NAD^+^ ratio (Licausi et al., 2010; Wagner et al., 2019). This affects downstream metabolic fluxes and thus forcing the system to undergo a metabolic shift (Bailey-Serres and Voesenek, 2008). Despite this energy crisis, basic molecular processes including transcription and translation proceed, albeit with modified rates, and are critical for acclimation (Fennoy et al., 1998; Branco-Price et al., 2005; Mustroph et al., 2009). This implies that protein import into the ER continues and that the lack of oxygen would lead to an accumulation of non-oxidized proteins in the ER. Our results show that diminished ERO increases the severity of this problem, likely by generating a bottleneck in the oxidation capacity of the lumen, which cannot be compensated by other less efficient oxygen-dependent systems independent of ERO. Under natural conditions such situations may frequently occur with water logging when particularly roots suffer severe oxygen deprivation. In this case, alternative pathways for electron dissipation from ERO FAD may help to ensure survival.

### Arabidopsis EROs are ER-resident type II membrane proteins

To fulfill their predicted molecular function in disulfide formation on proteins either resident in the ER lumen or passing through the ER for secretion, EROs need to be located in the early secretory pathway. This leads to the question of how permanent residency in the ER is achieved to ensure efficient oxidative protein folding. Consistent with biochemical studies on soybean ERO1a (Matsusaki et al., 2016), we have shown that both EROs of Arabidopsis are type II proteins with their N-termini facing the cytosol (Figure 3). The hydrophobic patch of 21 amino acids close to the N-terminus hence appears to function as targeting signal and membrane anchor simultaneously. A net positive charge among the N-terminal stretch of 13 amino acids preceding the TMD is likely key for orienting the protein with its N-terminus on the cytosolic side of the membrane according to the positive-inside rule (von Heijne, 1992; Lerch-Bader et al., 2008). This is fully consistent with multiple other type II proteins (Goder and Spiess, 2003). Expression of an N-terminal fragment of ERO1 with only 37 amino acids including the TMD as N- or C-terminal roGFP2 fusions also strongly suggests that this N-terminal domain is sufficient for correct ER targeting and orientation of the TMD (Supplemental Figure S5D). In the absence of any obvious retrieval signal, it is most likely the TMD itself that restricts EROs from leaving the ER with normal membrane flow. Shorter TMDs are generally considered to act in protein retention (Cosson et al., 2013), but other determinants within the TMD cannot be excluded at this point. Other solutions for ER retention have evolved in non-plant species: (*i*) human cells employ the formation of mixed disulfides with PDIs that are prevented from secretion by a C-terminal –KDEL or –RDEL retrieval motif (Anelli et al., 2002; Anelli et al., 2003; Otsu et al., 2006); (*ii*) In yeast, Ero1p is also a membrane protein albeit as a type I protein inserted with a C-terminal TMD (Pagani et al., 2001). Despite the different solutions to achieve retention, the unifying consequence is that ER residency of EROs is conserved.

The obtained results unambiguously show that a simplified ReTA assay with a merge of two redox-dependent fluorescent signals rather than a quantitative ratiometric analysis is sufficient to distinguish N- and C-terminal fusions of the respective protein with roGFP2 (Figure 3). However, in the case of fusion of roGFP2 to the ERO C-termini, that are in close proximity to the catalytic domain, the dynamic range of the probe was diminished by about 50% compared with soluble roGFP2 in the ER lumen (Figure 3, Supplemental Figure 5, (Brach et al., 2009). This is most likely due to the formation of mixed disulfides, which would severely compromise the use of such fusion proteins for dynamic redox measurements. This disadvantage cannot be circumvented, even if one would initially reduce the probe with DTT, as Merksamer et al. (2008) reported for application of roGFP2 in the ER of yeast cells.

### Grx1-roGFP2iL enables dynamic recording of glutathione redox potential in the ER lumen

Luminal pH and Ca^2+^ have been monitored in the ER of plants with GFP-based probes in the past (Bonza et al., 2013; Martinière et al., 2013; Shen et al., 2013; Resentini et al., 2021). When the most widely used redox-sensitive probe roGFP2 was expressed in the ER, it was found to be 99% oxidized due to the negative midpoint potential of the probe (-280 mV) and thus did not allow measurements of steady-state redox potentials with any degree of accuracy (-225 mV was calculated in tobacco epidermis assuming luminal pH at 7.2) nor dynamic physiological changes towards more oxidizing conditions (Schwarzländer et al., 2008). roGFP2 in the ER could only be used for dynamic measurements in the specialized context of severe reductive stress application pre-reducing luminal *E*_GSH_ by DTT (Merksamer et al., 2008). Strong oxidation of the probe in the lumen of the ER, however, was exploited for visualization of membrane protein topology (Figure 3; (Brach et al., 2009)). Here, expression of Grx1- roGFP2iL with a midpoint potential of -240 mV in the ER enabled fully dynamic measurements of *E*_GSH_ with responsiveness of the probe in both directions from the steady-state fluorescence ratio values and hence applicability of established calibration protocols. The steady-state *E*_GSH_ of -241 mV (Figure 4) is surprisingly reducing and at the reducing end of all values that have been reported for the ER of different non-plant cell types before (Schwarzländer et al., 2008; Kolossov et al., 2011; van Lith et al., 2011; Delic et al., 2012; Birk et al., 2013). Considering a concentration of 2.5 mM GSH in the cytosol (Meyer et al., 2007) and a passive concentration dependent transport of GSH into the ER, the determined steady-state *E*_GSH_ of -241 mV in wild-type cells according to the Nernst equation would imply a GSH:GSSG ratio of approximately 400:1. This is significantly less oxidizing than ratios between 1:1 and 3:1 suggested earlier (Hwang et al., 1992; Sevier et al., 2007). This contrasts a GSH:GSSG ratio of ∼50.000:1 in the cytosol where glutathione reductase continuously and efficiently reduces GSSG (Meyer et al., 2007; Marty et al., 2009; Schwarzländer et al., 2016).

Thiol redox potentials are sensitive to pH and sensor readout would need appropriate pH- adjustments for exact determination of absolute values (Schwarzländer et al., 2008). pH in the secretory pathway has been reported to decrease from near neutral values in the ER to increasingly acidic values beyond the Golgi (Shen et al., 2013; Schumacher, 2014) and thus the calibration protocol applied here with a theoretical titration curve for pH 7 appear valid for a reasonable approximation of the actual *E*_GSH_ in the ER.

ERO proteins introduce oxidation power by transferring electrons derived from thiols to molecular oxygen (Meyer et al., 2019). In the *ero1 ero2* plants generated in this work, diminished ERO activity shifted the *E*_GSH_ to a steady state value of -253 mV, which is consistent with a bottleneck in electron efflux from the luminal thiol-based redox systems. The shift of 12 mV towards more negative conditions does not affect the growth of the plants under non-stress situations. It does, however, have pronounced deleterious consequences under situations where the reductive load in a cell is increased by DTT exposure. Under those conditions of reductive ER stress, we recently found that mitochondrial respiration can act as a reductant sink for excess thiols. The capacity of this system is boosted under reductive stress through induction of alternative oxidase 1a (AOX1a). An *ero1 aox1a* double mutant showed synergistic sensitivity to DTT compared to the single mutants showing that the mitochondria contribute the safeguard the ER lumen from excess reductive burden (Fuchs et al., 2022).

### ERO activity mediates redox dynamics in the ER lumen

Environmental challenges typically cause deviations from homeostasis, which means that the biochemical steady state of many reactions is shifted but needs to be restored to prevent damage or inefficient usage of resources (Cramer et al., 2011; van Zelm et al., 2020). The ER lumen is characterized by far less reducing conditions with respect to its key Cys-based redox couples than in the cytosol, plastids, and mitochondria (Schwarzländer et al., 2008). DTT has long been used to cause protein folding stress in the ER through reductive challenge (Howell, 2013; Mishiba et al., 2019). The *in vivo* monitoring system that we introduce here now allows monitoring the effects of a transient reductive challenge and importantly the role of individual players, such as ERO, in the recovery (Figure 5). Diminished oxidizing power as a consequence of decreased ERO abundance also implies that in case of a sudden challenge, more time is required for re-adjusting metabolic pathways and steady states of metabolite pools. A net reductive challenge may also occur when the normal oxidizing power decreases. Severely restricted electron flux along the mitochondrial electron transport chain under hypoxia causes a backlog of electrons in the NAD_+_/NADH pool and thus effectively a reductive challenge (Wagner et al., 2019). Similarly, lack of oxygen as the terminal electron acceptor initially causes overreduction of the luminal glutathione pool (Figure 6), most likely because import of reducing equivalents persist. The reductive challenge, in this case, occurs primarily from continued import of nascent peptides into the ER, which can be stopped by blocking translation with CHX (Figure 7). The similarity of the reduction rates in wild-type and all analyzed *ero* mutants suggests that this response is independent of ERO activity. After a severe challenge, cells respond through transcriptional changes to organize a defense line against the challenge (Lee et al., 2011; Licausi et al., 2011; Schmidt et al., 2018), or they may respond through regulation of protein activities (Gibbs et al., 2011; Dissmeyer, 2019; Millar et al., 2019). The gradual return to more oxidizing conditions during the second phase of the hypoxia treatment and the lack of this response in *ero1 ero2* mutants (Figure 6) strongly suggests that EROs are the drivers of this oxidation. Only the residual activity in *ero1 ero2* appears insufficient to achieve a pronounced oxidation. The ERO1 promotor contains a UPR element rendering the gene responsive to protein folding stress in the ER (Supplemental Figure S3; (Martínez and Chrispeels, 2003; Fan et al., 2019). However, the maintenance of the re-oxidation response in *ero1-3* may indicate that ERO1 is not involved in this recovery, and that this gradual oxidation is not caused by *de novo* synthesis of oxidizing proteins during UPR. As an alternative, biochemical fine-tuning of EROs can be envisaged. Ero1p and mammalian Ero isoforms contain regulatory disulfides albeit in different positions on the protein (Appenzeller-Herzog et al., 2008; Baker et al., 2008; Sevier and Kaiser, 2008). Beyond the catalytic cysteines directly involved in electron transfer, plant ERO homologs also contain several additional cysteines that are highly conserved throughout the plant kingdom that may allow for posttranslational regulation of ERO activity (Aller and Meyer, 2013; Fan et al., 2019). To what extent such putative regulatory mechanisms are relevant *in vivo* remains to be shown. It remains unclear though, what the electron acceptor und the applied hypoxic conditions might be. The fact that the sensor still responds ratiometrically during re-oxygenation with sensing a transient pronounced oxidation before gradual recovery to values similar to starting values emphasizes that the cells are kept alive during the course of the experiment. With this, further dissection of this ERO-dependent oxidation in future work appears possible.

Besides nascent peptides, GSH is another reducing input in the ER lumen. Based on data from yeast (Ponsero et al., 2017), import of GSH into the ER is assumed to occur via the SEC61 translocon, even though this path has not been specifically confirmed in plants (Figure 8). Partial depletion of cellular GSH through preincubation of leaf discs in BSO for 18 h increased the fluorescence ratio of luminal Grx1-roGFP2iL (Figure 7C), indicating the luminal *E*_GSH_ to be less negative than in control leaves. The ER is thus not completely autonomous from the cytosol in its glutathione redox pool and follows the depletion of GSH in the cytosol. Increased luminal oxidation in GSH-depleted leaves highlights an important contribution of GSH in defining the redox potential in the ER. Depletion of GSH also led to far more pronounced oxidative recovery to values far beyond the original steady-state values and a delayed recovery towards the original values after re-oxygenation (Figure 7C, D). Although the underlying redox reactions are yet unknown, a similar response in wild type and *ero1 ero2* suggests that this process may be independent of ERO activity.

**Figure 8.**
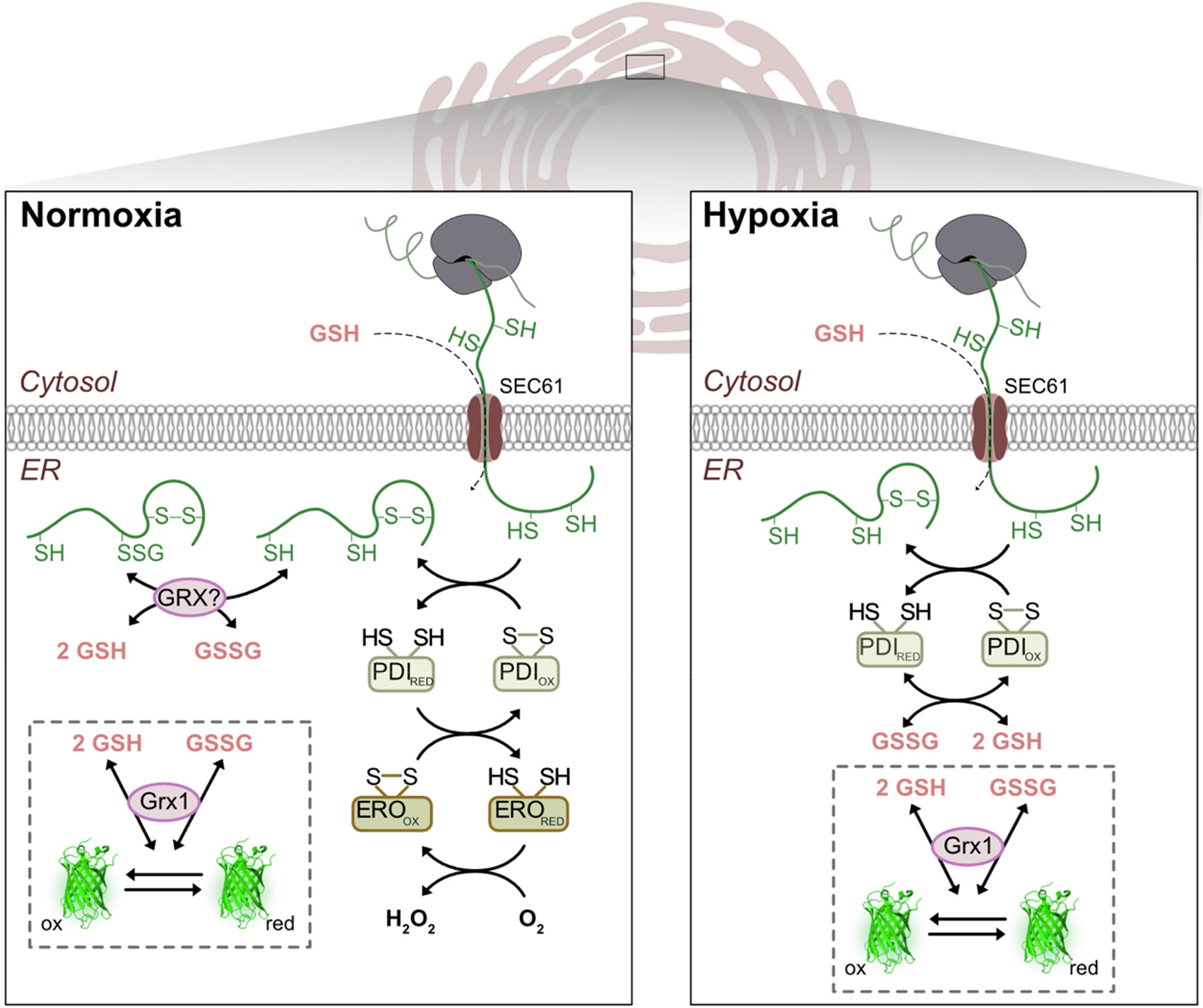
A model depicting redox homeostatic mechanism and thiol-disulfide exchange reactions in the ER under normoxia and hypoxia. Nascent peptides with reduced thiol residues and GSH enter the ER via the SEC61 translocon. The educing power is counteracted by oxidizing power generated by the PDI/ERO system that transfers electrons to molecular oxygen. The hypoxia model on the right depicts the early phase of hypoxia when still nascent peptides enter the ER while oxidative power that is normally generated by ERO is lost. Whether endogenous glutaredoxins participate in thiol-dithiol exchange reactions between glutathione and proteins remains unknown. The redox sensor Grx1-roGFP2iL is introduced into the ER lumen to read out the local *E*_GSH_.

### Conclusion

The results presented in this work demonstrate that ERO activity is key in establishing resilience against reductive stress and hypoxia. The obtained insight and technical development jointly raise several questions that deserve future attention. The experimental setup for dynamic redox studies in the ER of living plant cells is likely to be an important step towards a systematic dissection of the ER redox network and its key players (Figure 8). Such studies may involve 18 complementation with protein mutants to determine the function of specific residues for electron transfer reactions and null mutants for additional candidates involved in oxidative protein folding in the ER.

## MATERIALS AND METHODS

### Plant material and growth conditions

Arabidopsis (*A. thaliana* [L.] Heynh.) plants, ecotype Col-0, were grown on soil in growth chambers under long-day conditions (16 h light of approx. 120 μmol photons m^-2^ s^-1^ at 22°C, 8 h dark at 18°C) with a humidity of about 50%. Seeds were surface sterilized in 70% (v/v) ethanol for 5 min. Afterwards, seeds were washed twice in sterile water and transferred to Arabidopsis growth medium solidified with 0.8% (w/v) agar as described earlier (Meyer and Fricker, 2000). For preparation of plates supplemented with dithiothreitol (DTT), the autoclaved nutrient medium was cooled down to 50°C and sterile-filtered DTT was added to the desired final concentration shortly before gelling. Plates with seeds were stratified at 4°C for 24 hours before placing them in a controlled growth chamber in a vertical orientation under long-day conditions (16 h light approx. 75 μmol photons m^-2^ s^-1^ at 22°C, 8 h dark at 16°C). To test the sensitivity of seedlings to DTT, surface-sterilized seeds from wild-type and the indicated genotypes were germinated on Murashige & Skoog (MS) agar plates and after 5 days carefully transferred to plates supplemented with the indicated DTT concentration. Seedling growth was documented 5 days after transfer. Pictures of seedlings on plates were taken with a DSLR camera and root length measured with ImageJ (Schneider et al., 2012) (http://rsb.info.nih.gov/ij/). For treatment of Arabidopsis leaves with tunicamycin (TM) or DTT, leaf discs with a diameter of 5 mm without major veins were cut with a cork borer from 6-week old wild-type plants. Leaf discs were vacuum infiltrated with water supplemented with 5 µg/mL TM (in DMSO) or 2 mM DTT. As control, solution of 0.5% (v/v) DMSO or water were used, respectively. After vacuum infiltration, the samples were incubated for 6 hours.

### Screening of Arabidopsis T-DNA insertion lines

Seeds of wild-type (ecotype Columbia-0, Col-0) and the SALK T-DNA insertion lines SALK_004929 (*ero1-5*), SALK_003488 (*ero1-4*), and SALK_096805 (*ero1-3*) with a T-DNA insertion in the gene *ERO1* (At1g72280) and SALK_000573 (*ero2-3*) with a T-DNA insertion in the gene *ERO2* (At2g38960) were obtained through NASC. To isolate homozygous mutants, DNA was extracted from leaf material following established protocols (Edwards et al., 1991) and screened for T-DNA insertions in *ERO1* and *ERO2* by PCR using a T-DNA left border primer as well as the respective gene-specific primer pairs (Supplemental Table S1).

### RNA extraction and Semi-quantitative PCR

Plant tissue was disrupted mechanically using 3 mm tungsten carbide beads (Qiagen, Hilden, Germany), and a TissueLyser II (Quiagen) at 30 Hz for 2 min. RNA extraction was performed using NucleoSpin RNA isolation kit according to manufacturer’s protocol. 1 μg of total RNA was used for cDNA synthesis using the M-MLV reverse Transcriptase Kit (Invitrogen Ltd, Carlsbad, CA, USA) according to the manufacturer’s protocol. Instead of RNAseOut, the equal amount of RNase free deionized H_2_O was used. Semi-quantitative PCR was carried out on 1 μL of cDNA with gene-specific primers against *ERO1* (P7/P8), *ERO2* (P9/10) and *ACTIN7* (At5g09810; P11/P12).

### Cloning of 35S_pro:_amiR*ERO2*

The amiRNA targeted against *ERO2* (amiR*ERO2*) was designed using the MicroRNA Designer with the microRNA miR319a as a template to create the amiRNA hairpin (http://wmd3.weigelworld.org; Ossowski et al., 2008). Primer sequences are listed in Supplemental Table S1. The final amiRNA product was PCR-amplified with Gateway compatible overhangs flanking the sequence encoding the amiR*ERO2* hairpin (primer P23/P24). The resulting product was recombined into pDONR201 in the BP reaction. Positive clones were recombined into the plant binary vector pB2GW7 (Karimi et al., 2002). Transformation of *Arabidopsis* was performed by floral dip (Clough and Bent, 1998). Seeds were sown on soil and positive transformants were selected by application of 240 mg/L glufosinate-ammonium solution (Basta®; Bayer Crop Science, Monheim, Germany). Resistant plants were transferred to soil.

### Quantitative real-time-PCR

Total RNA was extracted leaf tissue using NucleoSpin RNAII Kit according to manufacturer’s protocol (Machery-Nagel, Düren, Germany). 1 μg of total RNA was used for cDNA synthesis using the M-MLV reverse Transcriptase Kit (Invitrogen) and an oligo(dT) primer. The PCR reaction was carried out with the MESA Green qPCR MasterMix Plus, no ROX using SYBRGreen as a dye in a 384 well plate on a CFX96 cycler (BioRad, Hercules, CA). The total volume of the reaction mix was 8 μL containing 250 μM of the 1:1 pre-mixed primer, 1 μL cDNA and 1x MESA Premix. To check the primer efficiency, a mixture of all cDNAs was used to plot a linear regression (1:2; 1:4; 1:8; 1:16; 1:32; 1:64; 1:128) of log(*n*) (*n* = number of template molecules) against the Ct value of the respective primer pair. PCR for each of the three biological replicates was performed in triplicates. The initial denaturing time was 5 min, followed by 54 cycles of 95°C for 15 s and 60°C for 60 s. A melting curve was run after the PCR cycles. Transcript abundance was measured with specific primers against *ERO1* (P25/P26), *ERO2* (P27/P28), and *SAND family protein* (At2g28390, P29/30).

### roGFP2 fusion constructs and transient transfection of *Nicotiana benthamiana* leaf cells

To obtain N- and C-terminal fusions of full-length ERO1 and ERO2 with roGFP2, the coding sequence of both ERO1 and ERO2 was amplified using Gateway-compatible primers. To allow C-terminal fusion of roGFP2 to ERO1 and ERO2, both genes were amplified with primers P13/14 and P16/P17, respectively. For N-terminal fusion to roGFP2, the ERO sequences were amplified using the primer pairs P13/P15 and P16/P18, respectively (Supplementary Table S1). The resulting fragments were purified and mixed with pDONR201 for the BP reaction. Positive clones were recombined in the LR reaction with the destination vector pSS01 to generate C-terminally tagged ERO1/2-roGFP2, or with the destination vector pCM01 to generate N-terminally tagged roGFP2-ERO1/2 fusion proteins respectively (Brach et al., 2009). For transient expression of the ERO fusion proteins, *N. benthamiana* Domin plants were grown in a growth chamber under controlled conditions and leaf epidermal cells transformed as described previously (Sparkes et al., 2006). For transformation, leaves were infiltrated with sterile deionized water containing *Agrobacterium tumefaciens* strain AGL1 harboring the respective binary vectors for expression of the roGFP2 fusion proteins. Transfected cells were imaged by confocal laser scanning microscopy three days after inoculation.

### Targeting of Grx1-roGFP2iL to the ER lumen and stable transformation of *Arabidopsis thaliana*

For ER targeting, *Grx1-roGFP2iL* (Aller et al., 2013) was PCR-amplified using the primers P19 and P20 to introduce *Nco*I and *Xba*I restriction sites. The amplified product was blunt end-ligated into pJet1.2 (http://www.thermoscientificbio.com) and confirmed through sequencing. Subsequently, the sensor was cloned as a *Nco*I/*Xba*I fragment into pWEN81 between a chitinase signal peptide and the HDEL retrieval signal. Afterwards, the entire cassette consisting of *Grx1-roGFP2iL* with the N-terminal chitinase ER-targeting signal and the C-terminal HDEL was PCR amplified using the primers P21 and P22 and sub-cloned into the binary vector pBinCM under the control of a constitutive *UBIQUITIN10* promoter (At4g05320*; UBQ10*; (Grefen et al., 2010) using *Kpn*I and *Sal*I restriction sites. The resulting construct was transformed into *A*. *tumefaciens* strain AGL1. Arabidopsis plants were transformed with Agrobacterium through floral dip (Clough and Bent, 1998). Positive transformants were visually screened for GFP fluorescence on a stereomicroscope (Leica M165FC, Leica, Wetzlar, Germany) and transferred to soil for seed production.

### Confocal laser scanning microscopy

Confocal imaging was carried out on a Zeiss confocal microscope LSM780 (Carl Zeiss Micro Imaging, Jena, Germany) equipped with lasers for 405, 488 and 543 nm excitation. Images were collected with a 40x lens (C-Apochromat 40x/1.2 W Corr M27, Zeiss) in a multi-track mode with line switching between 488 nm and 405 nm excitation and taking an average of four readings. The fluorescence emission was collected at 505–530 nm for roGFP2 and at 590–630 for mCherry. Autofluorescence excited at 405 nm was collected for the band 430–470 nm and values were used to subtract autofluorescence bleeding into the roGFP channel as described previously (Schwarzländer et al., 2008; Fricker, 2016). For sequential perfusion of different treatments, 4–5-day old seedlings were mounted in a RC-22 perfusion chamber mounted on a P1 platform using a steel anchor harp with a 1.5 mm grid mesh (Warner Instruments, Hamden CT). Imaging buffer, and the indicated solutions of DTT and H_2_O_2_, were loaded into 50 mL syringes connected to a VC-8M valve controller (Warner Instruments) with 1.5 mm polyethylene tubes (Ugalde et al., 2020).

### Ratiometric analysis of confocal images

Images were imported into a custom written MatLab (The MathWorks, www.mathworks.de) analysis suite version v1.3 (Fricker, 2016). The ratio analysis was performed on a pixel-by-pixel basis as *I*405/*I*488 following spatial averaging in (*x*,*y*) using a 3 × 3 kernel. Correction of *I*405 for autofluorescence bleeding into the 405 nm channel and subtraction of background signals for each channel was performed. The average background signal was typically measured from the vacuole of one of the cells. For pseudo-color display, the ratio was coded by hue on a spectral color scale ranging from blue (fully reduced) to red (fully oxidized), with limits set by the *in situ* calibration. Images were collected close to the bottom of epidermal cells to identify the nucleus with its ER-characteristic nuclear ring. Due to its midpoint redox potential of -280 mV (Dooley et al., 2004), roGFP2 located in the ER lumen is almost completely oxidized. In contrast, roGFP2 located in the cytosol is almost completely reduced (Meyer et al., 2007). For Arabidopsis seedlings stably expressing Grx1-roGFP2iL-HDEL in the ER, the calibration was done by pre-incubation in 10 mM DTT and 10 mM H_2_O_2_ to drive the sensor to its fully reduced and fully oxidized form, respectively. Ratio values were log_10_ transformed before plotting to convert skewed ratio data distribution with unequal variance to a normal distribution.

### Determination of the degree of oxidation

The degree of sensor oxidation (*OxD*) for the Grx1-roGFP2iL-HDEL sensor was determined using the following equation:

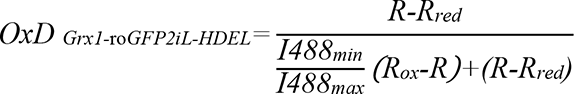

for which *R*, *R_red_* and *R_ox_* denote the 405/488 ratio of the fluorescence intensities for excitation at 405 nm and 488 nm fluorescence ratios at steady-state (*R*), upon full reduction (*R_red_*) or oxidation (*R_ox_*), respectively. *I488_min_* and *I488_max_* represent the fluorescence intensities of fully oxidized and fully reduced roGFP2iL. The calculation of *OxD* was done from the mean fluorescence data. Titration curves drawn for Grx1-roGFP2iL were calculated from the Nernst-Equation with *E_0_’_roGFP2iL_*= -240 mV.

### Hypoxia treatments

Seedlings of WT and *ero1 ero2* were grown on vertically oriented square plates with 0.5x MS supplemented with 0.5 (w/v) sucrose. 12 days after germination, plates with 12-15 seedlings where exposed to anoxia (0% O_2_) in the dark such that gradually remaining O_2_ could diffuse out of the plates. When the O_2_ concentration reached 0.3%, seedlings were kept under these conditions for 8.5 hours. Subsequently, plates were returned to long-day conditions (16 hours light, 8 hours dark) and kept for another 3 days. For evaluation of hypoxia effects, a survival score was determined as described earlier (Gibbs et al., 2011). Briefly, plants were categorized as healthy (5 points), damaged (3 points) and dead (1 point). The number of plants in a category was multiplied by the respective score, summed up and divided be the total number of seedlings in the experiment to get the final survival score. To test the effect of hypoxia on soil grown plants at rosette stage, 5-week-old plants were grown under short-day conditions and then transferred to black boxes in which they were flooded with tap water to a level 10 cm above the rosette. Boxes were covered to prevent all photosynthetic activity. Control plants were kept under normal short-day conditions. After 4 or 5 days of dark-submergence, plants were taken out of the boxes and maintained under short-day conditions. On day 6 plants were individually photographed and analyzed for their fresh weight.

To test the effect of hypoxia on primary root growth, five-day old seedlings from the indicated genotypes were placed for 3 h in a 2.5 L AnaeroGen^TM^ chamber equipped with an Oxoid^TM^ bag to generate hypoxic conditions (Thermo Scientific, Waltham, MA). After the treatment, plates were kept for 48 h under normal normoxic growth conditions and documented at the end.

For time-resolved ratiometric analysis of the Grx1-roGFP2iL sensor under hypoxia, leaf disks from 4–6-week-old plants of the indicated genotypes were placed in a 96-well plate and submerged in imaging buffer. Leaf discs of 7 mm diameter were cut out with a cork borer avoiding the major veins. Sensor fluorescence was measured using a CLARIOstar plate reader equipped with an atmospheric control unit (ACU) (BMG Labtech, Ortenberg, Germany). Hypoxic conditions (0.1% O_2_) were reached by automatically pumping N_2_ gas into the plate reader chamber controlled by the ACU system. roGFP2 was collected at 520 ± 5 nm, after subsequent excitation by a filter-based excitation system at 400 ± 5 nm and 480 ± 5 nm. Orbital averaging of fluorescence readings along a 3 mm-diameter circle was used to account for heterogeneous distribution of signal across the wells, according to (Ugalde et al., 2021). Autofluorescence was estimated separately for each genotype by measuring the fluorescence in leaf discs taken from the respective non-transformed plants. These values were subtracted from the readings taken for roGFP2iL reporter lines. A custom Python script was used to subtract the fluorescence and calculate the 400/480-nm ratio for each sample. Ratio values were log_10_ transformed before plotting. For pharmacological treatments, 70 µM cycloheximide (CHX), and 1 mM L-Buthionine-(S,R)-sulfoximine (BSO), respectively, were added to each well and leaf discs incubated in the dark for 18 h before the measurements. All samples remain in the same solutions during the subsequent 8 h measurement period.

## Accession numbers

Sequence information can be found in the GenBank repository under these accession numbers: *ERO1*, At1g72280; *ERO2*, At2g38960; *SAND* family protein, At2g28390; *UBIQUITIN10,* At4g05320.

## Author contributions

A.J.M., I.A., S.J.M.-S. and M.Schw. designed the research; J.M.U., I.A., R.R.S., M.H., P.F., S.L., L.K., M.Schl. and M.Schw. performed the experiments and analyzed results; J.M.U., M.Schw. and A.J.M. wrote the manuscript with input from all other authors.

## SUPPLEMENTAL DATA

Supplemental Figure S1. Analysis of segregating T-DNA insertion lines for *ERO1* and *ERO2*.

Supplemental Figure S2. Generation of viable *ero1 ero2* double mutants.

Supplemental Figure S3. E*R*O1 expression is induced under ER stress.

Supplemental Figure S4. e*r*o1 *ero2* seedlings are hypersensitive to reductive stress.

Supplemental Figure S5. Localization and orientation of ERO1 and ERO2 through ratiometric imaging of roGFP2 fusions.

Supplemental Figure S6. Generation of Grx1-roGFP2iL-HDEL for measurement of the glutathione redox potential in the ER.

Supplemental Figure S7. The ER-lumen of *ero1 ero2* is less oxidizing than in WT or single *ero* mutants.

Supplemental Figure S8. Hypoxia-induced changes in ER redox homeostasis can be monitored with Grx1-roGFP2iL.

Supplemental Figure S9. E_G_SH in the ER reflects the equilibration of reducing and oxidizing inputs.

Supplemental Table S1. Primers used in this study.

