## Supplemental data for "Endoplasmic reticulum oxidoreductin (ERO) provides resilience against reductive stress and hypoxic conditions by mediating luminal redox dynamics"

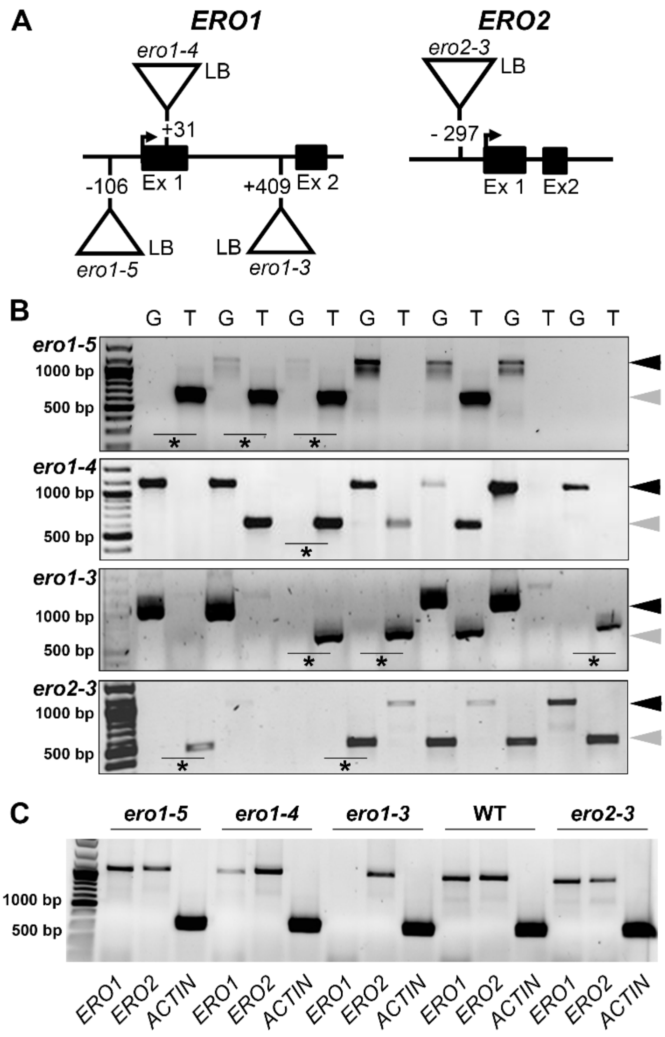


**Supplemental Figure S1. Analysis of segregating T-DNA insertion lines for *ERO1* and *ERO2*.** A, Gene models for the first two exons (Ex) of *ERO1* and *ERO2*. Insertion sites for different T-DNA lines are shown as triangles along the genes (*ero1-4*, SALK_003488; *ero1-5*, SALK_004929; *ero1-3*, SALK_096805 and *ero2-3* SALK_000573. The orientation of the T-DNAs is indicated by their left borders (LB). B, Homozygous lines for the indicated insertion lines were isolated from segregating populations via PCR using either two gene-specific primers (G, black arrow heads) or a combination of a gene-specific primer and a T-DNA primer (T, grey arrow head). Homozygous lines are indicated in the gel (*). C, Semi-quantitative analysis of *ERO1* and *ERO2* expression in the indicated homozygous T-DNA insertion lines. *ACTIN7* (AT5G09810) was used as a reference gene. For the sequences of all primers, see Supplemental Table S1.

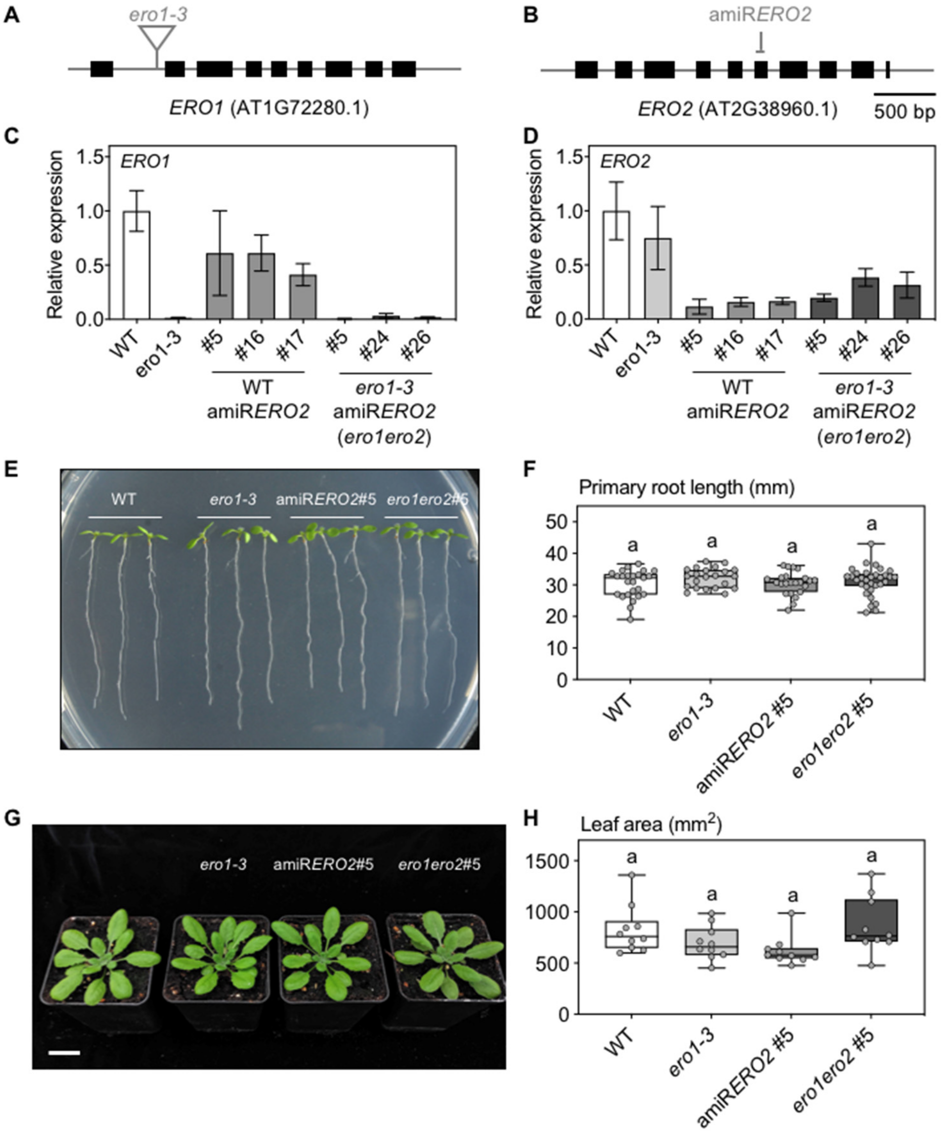


**Supplemental Figure S2. Generation of viable *ero1 ero2* double mutants.** A-B, Gene models for *ERO1* (A) and *ERO2* (B) indicating the insertion point for the T-DNA in *ero1-3* (SALK_096805) and the target sequence for the artificial microRNA, amiR*ERO2*. C, qPCR analysis of *ERO1* and *ERO2* transcripts in WT, *ero1-3* and lines expressing amiR*ERO2* in the WT (lines #5, #16 and #17) or in *ero1-3* (lines #5, #24 and #26). Bars indicate the mean transcript accumulation ± SD relative to the *SAND* housekeeping gene (At2g28390), *n* = 2–10. E, Representative image of 1-week-old WT, *ero1-3*, amiR*ERO2*#5 and *ero1 ero2*#5 seedlings grown on MS media. Bar = 1 cm. F, Primary root lengths for the genotypes depicted in panel E. *n*= 22–25. G, Representative images of 4-week-old WT, *ero1-3*, amiR*ERO2*#5 and *ero1 ero2*#5 plants grown on soil. Bar = 2 cm. H, Rosette leaf area for the genotypes depicted in panel G. *n*= 9–10. Although occasionally dwarf phenotypes were observed, this phenotype was not stable and lines could be maintained over several generations. This indicates that a minimum expression of *ERO2* is required in the *ero1 ero2* double mutant to maintain viability. For box plots in panels F and H, Box = interquartile range between the lower and upper quartiles, center line = median, whiskers = min and max values. Statistical analyses for data in panels F and H were performed using ANOVA with Fisher's LSD test. The letters indicate primary root length and leaf area in different genotypes were not statistically different.

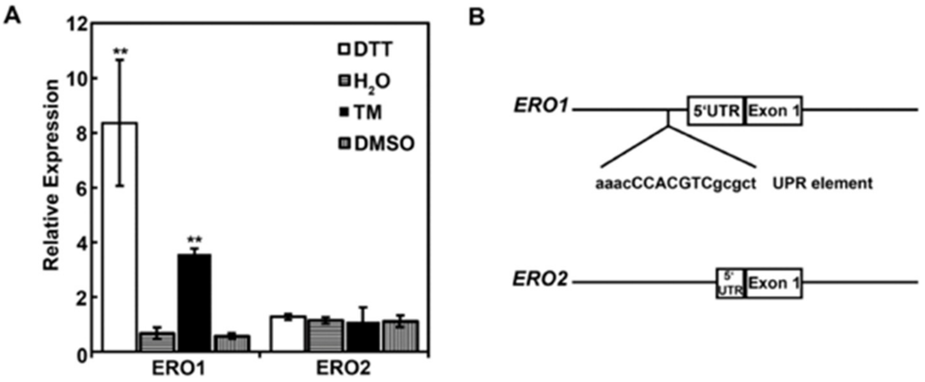


**Supplemental Figure S3. *ERO1* expression is induced under ER stress**. A, qPCR analysis of *ERO1* and *ERO2* transcripts in WT leaf disks treated for 6 h with 2 mM DTT or 5 µg/mL tunicamycin (TM). Solvent controls were made by incubating samples for the same time in deionized water, or 0.5% (v/v) DMSO. Bar charts indicate the mean relative expression ± SD, $n$ = 3. Two-tailed Student's t-test was performed against the respective water and DMSO controls for DTT and TM, respectively, with ** $P$<0.01. B, Promoter regions of *ERO1* and *ERO2* showing only the promoter of ERO1 contains a classical unfolded protein response (UPR) element upstream of the 5' UTR region.

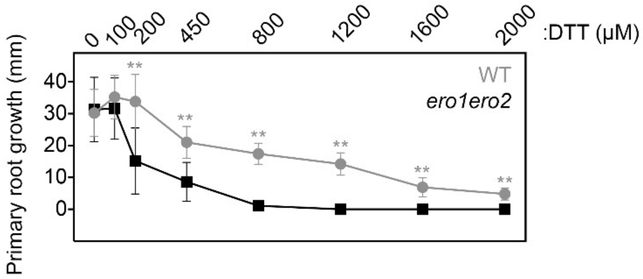


**Supplemental Figure S4. *ero1 ero2* seedlings are hypersensitive to reductive stress.** Primary root growth of WT and *ero1 ero2* seedlings after transfer to an extended range of DTT concentrations. Data indicate mean growth of primary roots ± SD, *n* = 18–42. Statistical differences were estimated after an unpaired t-test analysis. ** (*P*<0.05).

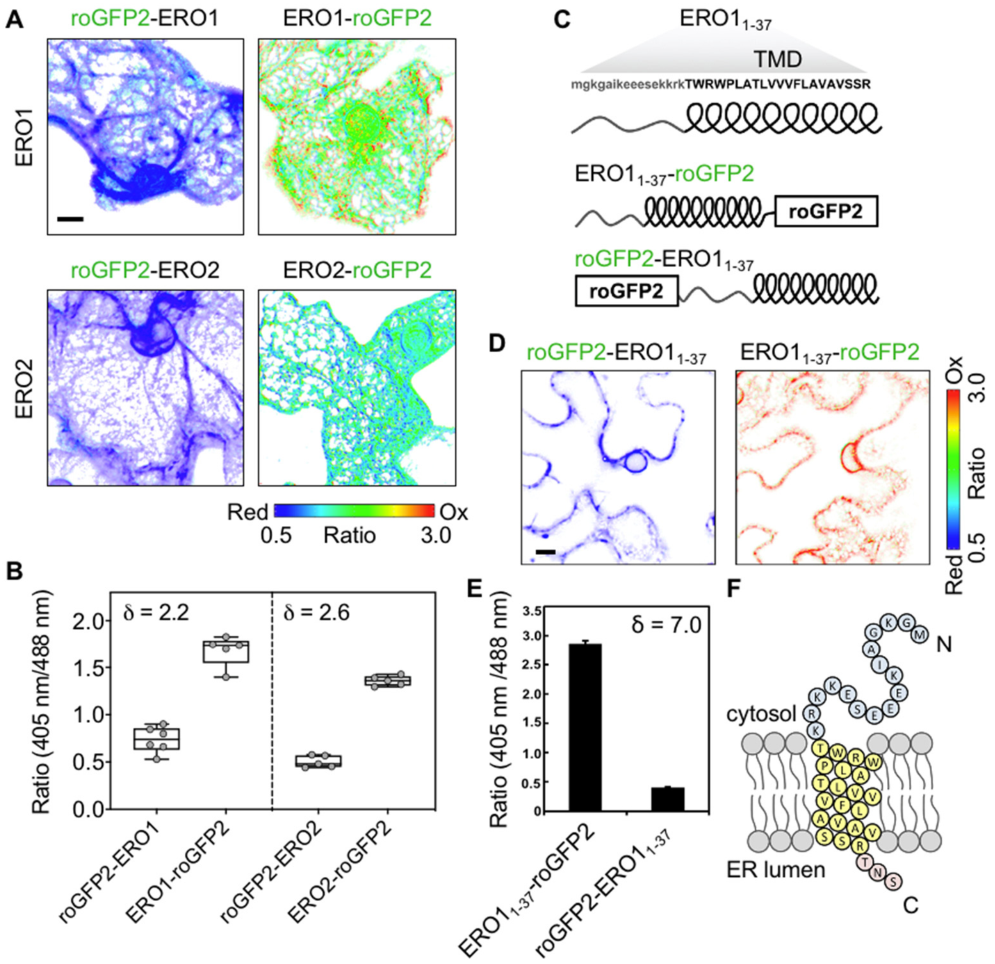


**Supplemental Figure S5. Localization and orientation of ERO1 and ERO2 through ratiometric imaging of roGFP2 fusions.** A, Representative ratiometric images of *Nicotiana bethamiana* leaf epidermal cells transiently expressing the roGFP2 sensor fused to either the N- or the C-termini of ERO1 (top panels) or ERO2 (bottom panels). Scale bar = 10 µm. False colors indicate the fluorescence ratios between the individual channels for roGFP2 (1: $\lambda_{ex}$ = 405 nm, $\lambda_{em}$ = 508–530 nm; 2: $\lambda_{ex}$ = 488 nm, $\lambda_{em}$ = 508–530 nm) on a scale from blue (reduced) to red (oxidized) defined by N- and C-terminally tagged SEC22 constructs (Brach et al., 2009). B, Fluorescence ratios for the roGFP2 fusions shown in panel A. *n* = 5. C, Scheme of the first 37 amino acids from the N-terminus of ERO1 containing the predicted transmembrane domain (TMD; capital black letters) (top panel). Middle and bottom panels show the fusion of roGFP2 to the C- or N- termini of ERO1$_{1-37}$, respectively. D, Representative ratiometric images of *N. benthamiana* leaf epidermal cells transiently expressing roGFP2-ERO1$_{1-37}$ or ERO1$_{1-37}$-roGFP2 fusions. Despite the consistent readout for the orientation of EROs, quantitative analysis of the roGFP2 ratios revealed that at steady-state ERO1/2-roGFP2 did adopt lower ratios than expected for full oxidation. The actual ratio changes between N- and C-terminal fusions respective dynamic range of roGFP2 in this case was 2.2-2.6 (B). Because these values are much lower than the full dynamic range (δ) of 5.5 observed in prior work (Brach et al., 2009), we concluded that EROs may directly interact with roGFP2, likely though formation of mixed disulfides when they are forced into close proximity in fusion proteins. When only a 37-amino acid N-terminal fragment of ERO1 that included the TMD was used with either N- or C-terminal roGFP2 a larger δ reflecting the full redox gradient across the ER membrane was found (C-F). Scale bar = 10 µm. E, Fluorescence ratios calculated for the fusion proteins shown in panel D. *n* = 5. F, Scheme of the TMD within the first 37 amino acids of the N-terminal from ERO1. For box plots: Box = interquartile range between the lower and upper quartiles, center line = median, whiskers = min and max values.

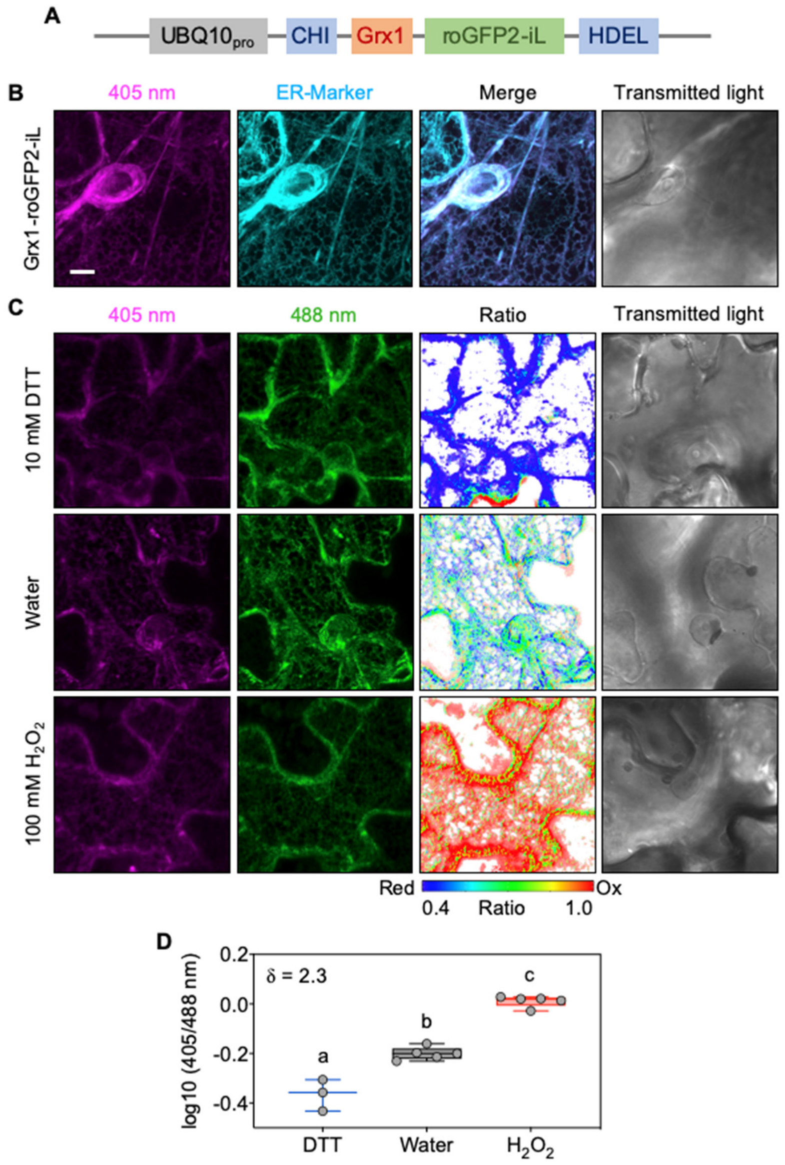


**Supplemental Figure S6. Generation of Grx1-roGFP2iL-HDEL for measurement of the glutathione redox potential in the ER**. A, Map of the sensor construct for ER-targeted Grx1-roGFP2iL. Grx1-roGFP2iL (Aller et al., 2013) was cloned behind the CHITINASE target peptide (CHI) and tagged at its C-terminus with the ER retrieval signal HDEL. The construct was constitutively expressed under the control the Ubiquitin 10 promoter from Arabidopsis (*UBQ10$_{pro}$*). B, Confocal microscopy images of tobacco leaf epidermal cells transiently expressing Grx1-roGFP2iL-HDEL and the ER marker AtWAK2$_{TP}$-mCherry-HDEL (Nelson et al., 2007). roGFP2iL fluorescence was collected at 505–530 nm after excitation at 405 nm while the ER-marker was excited at 543 nm and fluorescence collected at 590–630 nm. Co-localization is depicted in the merged image. C, Responsiveness of Grx1-roGFP2iL in the ER of tobacco cells. For full reduction or full oxidation, tobacco leaf disks were vacuum infiltrated with either 10 mM DTT or 100 mM $H_2O_2$, respectively, or with deionized water as control. Dual excitation at 405 nm and 488 nm and emitted light collected at 505–530 nm enabled the calculation of 405 nm/488 nm ratio images. False colors indicate the fluorescence ratios on a scale from blue (reduced) to red (oxidized). Scale bar = 10 µm. D, Fluorescence ratios for the images shown in panel C after the indicated treatments, $n$ = 3–5. δ = dynamic range of the sensor. Box = interquartile range between the lower and upper quartiles, center line = median, whiskers = min and max values. Statistical analyses were performed using ANOVA with Fisher's LSD test. Different letters indicate statistically different groups ($P<0.05$).

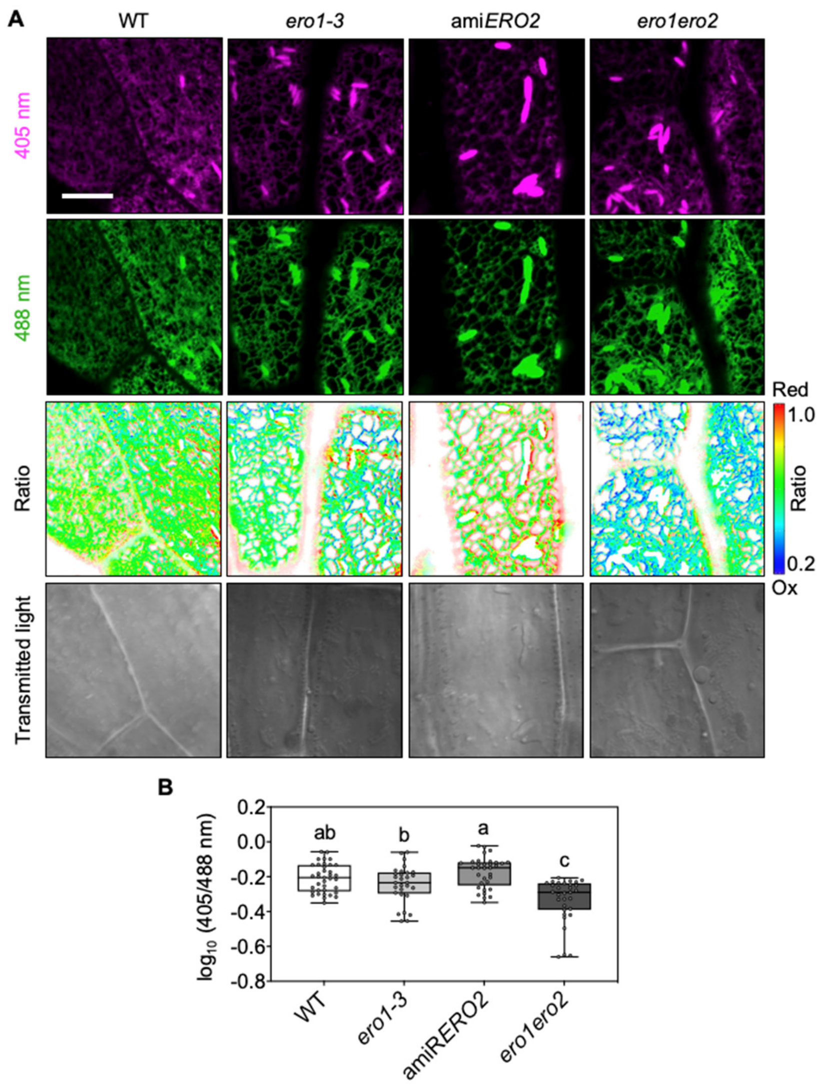


**Supplemental Figure S7. The ER-lumen of *ero1 ero2* is less oxidizing than in WT or single *ero* mutants.** A, Representative confocal images of hypocotyl cells of five-day old Arabidopsis seedling stably expressing Grx1-roGFP2iL-HDEL in the ER. roGFP2iL fluorescence was collected at 505–530 nm after successive excitation with 405 nm and 488 nm. Ratio images were calculated as the 405 nm/488 nm fluorescence. False colors indicate the fluorescence ratio values on a scale from blue (reduced) to red (oxidized). Scale bar = 10 µm. B, Fluorescence ratios for the samples shown in panel A, $n = 29$-$37$. Box = interquartile range between the lower and upper quartiles, center line = median, whiskers = min and max values. Statistical analyses were performed using ANOVA with Fisher's LSD test. Different letters indicate statistically different groups ($P<0.05$). Images for WT and *ero1 ero2* are the same as in Figure 4.

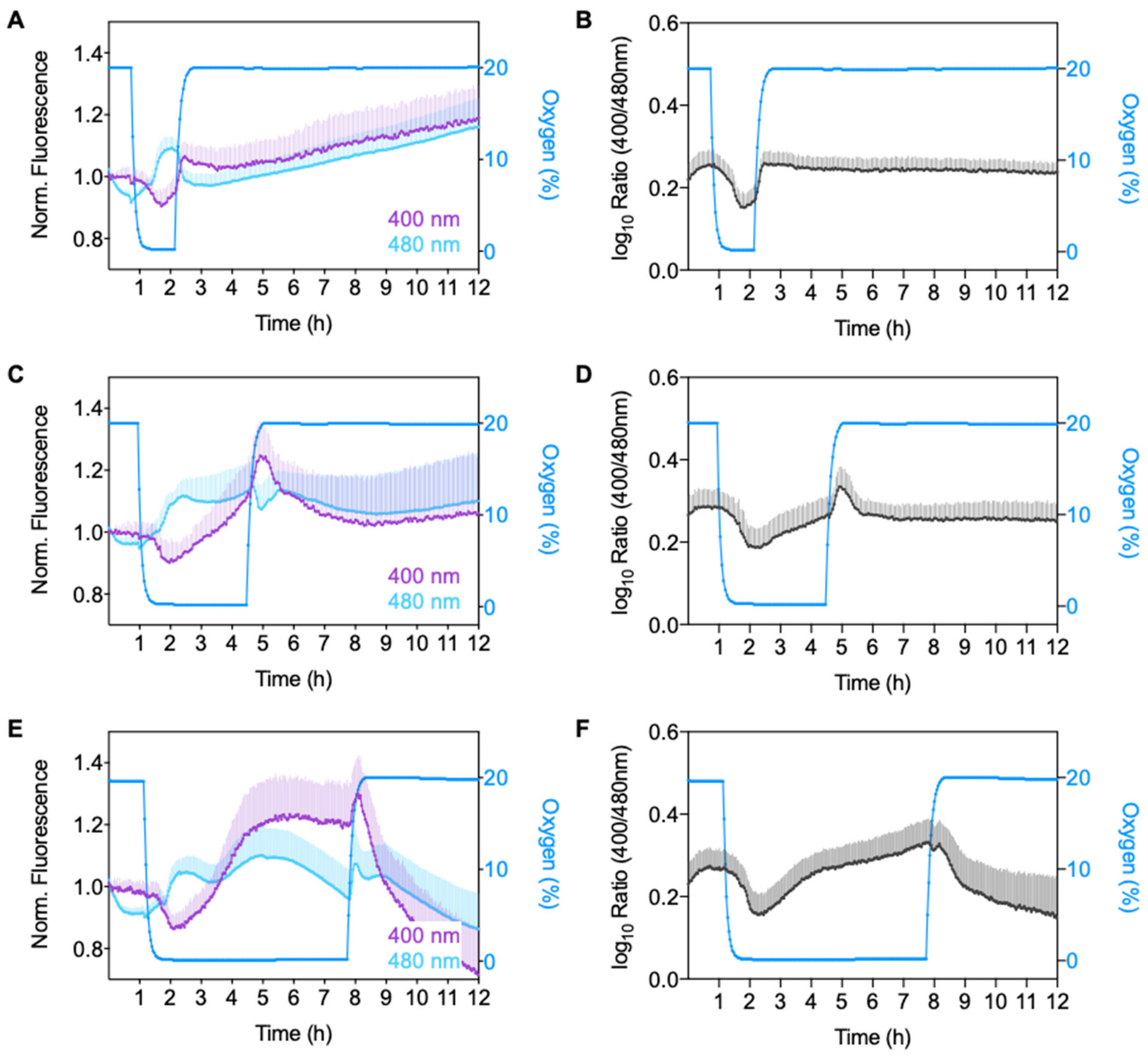


**Supplemental Figure S8. Hypoxia-induced changes in ER redox homeostasis can be monitored with Grx1-roGFP2iL.** A-C, Leaf disks from four-week-old wild-type plants expressing Grx1-roGFP2iL-HDEL were exposed to different oxygen conditions inside a plate reader equipped with an atmospheric control unit. roGFP2iL fluorescence was continuously collected at 520 ± 10 nm after excitation at 400 ± 5 nm and 480 ± 5 nm. Sensor fluorescence in leaf disks was measured initially for 1 h at 20% $O_2$ (normoxia), before $O_2$ was decreased to 0.1% (hypoxia) for 1 h (A), 3.5 h (B) or 6.5 h (C). At the end of the hypoxic phase normal $O_2$ levels of 20% were restored and fluorescence recorded for several hours to complete a full 12-hour time course. It was noted that during the extended hypoxic phases the two fluorescence excitation channels did not always show the expected opposing trends of a characteristic sensor response. Instead, the increasing ratio values under extended hypoxic conditions were primarily caused by selective increase of fluorescence excited at 400 nm. Because the analysis protocol includes subtraction of autofluorescence measured for non-transformed wild-type control samples on the same plate, we exclude the possibility of stress-induced accumulation of autofluorescent metabolites like e.g. anthocyanins being the cause for this increase. Despite this yet unexplained sensor response, the sensor retained its capability for dynamic redox equilibration, which is apparent by a bonafide ratiometric response during re-oxygenation after a preceding 3.5-hour hypoxic phase.

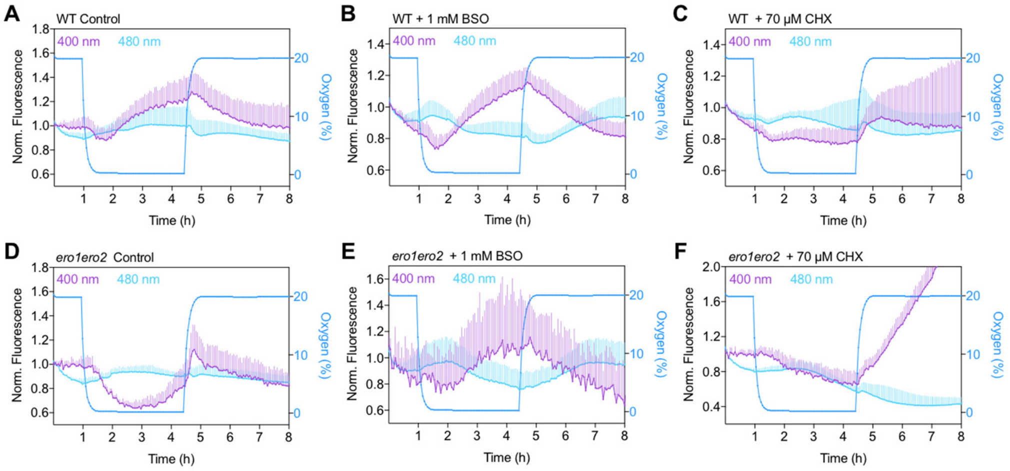


**Supplemental Figure S9. $E_{GSH}$ in the ER reflects the equilibration of reducing and oxidizing inputs.** Effect of cycloheximide (CHX, 70 µM) and buthionine sulfoximine (BSO, 1 mM) on hypoxia-induced redox dynamics in the ER of WT and *ero1 ero2* leaf disks. In all cases, leaf disks immersed in imaging buffer were used as controls. The data show the mean + SD of the normalized fluorescence for the independent 400 ± 5 nm and 480 ± 5 nm excitation channels for the effect of CHX (C and F), BSO (B and E), and imaging buffer as control (A and D) used to calculate the ratios presented in Figure 7. *n* = 4–9.

**Supplemental Table S1.** Primers used in this study

Primers used in semi-quantitative RT-PCR

| | |
|---|---|
| P1 | ATGGGAAAAGGCGCAATCAAA |
| P2 | CGCAATCCATTAGTGCACTTATATTTCTGAAT |
| P3 | ATGGCGGAGACGGACGTC |
| P4 | ACAGTCCATTATTGCACTTATGTTTCTGAATTG |
| P5 | AACCTCAGGACAACGGAATCTC |
| P6 | CAACCGGTATTGTGCTCGATTC |

Primers used to generate roGFP fusions with full length *At*ERO1 and *At*ERO2

| | |
|---|---|
| P7 | ggggacaagtttgtacaaaaaagcaggcttcATGGGAAAAGGCGCAATCAAA |
| P8 | ggggaccactttgtacaagaaagctgggtcCCAGAATGAGACAGCTAAATCC |
| P9 | ggggaccactttgtacaagaaagctgggtcTCACCAGAATGAGACAGCTAAATCC |
| P10 | ggggacaagtttgtacaaaaaagcaggcttcATGGCGGAGACGGACGTC |
| P11 | ggggaccactttgtacaagaaagctgggtcGCTTCTCTTCCCAGATACAGC |
| P12 | ggggaccactttgtacaagaaagctgggtcCTAGCTTCTCTTCCCAGATACAGC |

Primers used for cloning Grx1-roGFP2iL that is localized
in the lumen of the Endoplasmic Reticulum

| | |
|---|---|
| P13 | ACCATGGTGAGCAAGGGCGAGGAG |
| P14 | TCTAGACTTGTACAGCTCGTCCAT |
| P15 | AGGTACCATGGCTCAAGAGTTTGTGAAC |
| P16 | TATGTCGACTTACTTGTACAGCTCGTCCAT |

Primers used to generate ami*RERO2*

| | |
|---|---|
| I-2miR | gaTAGTATAGGTTCTGAACGCGAtctctcttttgtattcc |
| II-2miR | gaTCGCGTTCAGAACCTATACTAtcaaagagaatcaatga |
| III-2miR | gaTCACGTTCAGAACGTATACTTtcacaggtcgtgatatg |
| IV-2miR | gaAAGTATACGTTCTGAACGTGAtctacatatatattcct |
| P17 | ggggacaagtttgtacaaaaaagcaggcttcCTGCAAGGCGATTAAGTTGGGTAAC |
| P18 | ggggaccactttgtacaagaaagctgggtcGCGGATAACAATTTCACACAGGAAACAG |

Primers used in quantitative real-time PCR

| | |
|---|---|
| P19 | GACACAGATAGTGGTGAGATGAG |
| P20 | GGACAGTTCTCTGAATATATGGAG |
| P21 | ACTGACAATGATGAAATGACATATG |
| P22 | TTGGCAAGATCCTTCAGATGTAT |
| P23 | CCATATTGCAAGAAGTTTGCGCGTCTG |
| P24 | GCAAGTCATCGGGATGGAGAGACG |

Primers used for genotyping of T-DNA insertion lines

| | |
|---|---|
| #1393 | TCAAGAGCCAAAGATGAAACC |
| #1394 | AAATCAGTGGCACATTTCAGG |
| #1395 | ATTGTTTCTGGCCACAATTTG |
| #1396 | CCCTGAATCCATTGGCTAAAC |
| #1397 | AAATCAGTGGCACATTTCAGG |
| #1398 | TCAAGAGCCAAAGATGAAACC |
| #1399 | CCTGAGGTTTTCCCTCTTGAC |
| #1400 | CACAACAAACACAACAATGGG |
| #1401 | ATTTTGCCGATTTCGGAAC |
| #2726 | AAGGAAGACCTGGGACAAATG |
| #2727 | GCGTGTTATACAAAACCCACG |
| #2728 | GGTTTTGAGAAATTCCGGAAG |
